# Single-cell mapping of regulatory DNA:Protein interactions

**DOI:** 10.1101/2024.12.31.630903

**Authors:** Wei-Yu Chi, Sang-Ho Yoon, Levan Mekerishvili, Saravanan Ganesan, Catherine Potenski, Franco Izzo, Dan Landau, Ivan Raimondi

## Abstract

Gene expression is coordinated by a multitude of transcription factors (TFs), whose binding to the genome is directed through multiple interconnected epigenetic signals, including chromatin accessibility and histone modifications. These complex networks have been shown to be disrupted during aging, disease, and cancer. However, profiling these networks across diverse cell types and states has been limited due to the technical constraints of existing methods for mapping DNA:Protein interactions in single cells. As a result, a critical gap remains in understanding where TFs or other chromatin remodelers bind to DNA and how these interactions are perturbed in pathological contexts. To address this challenge, we developed a transformative single-cell immunotethering DNA:Protein mapping technology. By coupling a species-specific antibody-binding nanobody to a cytosine base editing enzyme, this approach enables profiling of even weak or transient factor binding to DNA, a task that was previously unachievable in single cells. Thus, our Docking & Deamination followed by sequencing (D&D-seq) technique induces cytosine-to-uracil edits in genomic regions bound by the target protein, offering a novel means to capture DNA:Protein interactions with unprecedented resolution. Importantly, this technique can be seamlessly incorporated into common single-cell multiomics workflows, enabling multimodal analysis of gene regulation in single cells. We tested the ability of D&D-seq to record TF binding both in bulk and at the single-cell level by profiling CTCF and GATA family members, obtaining high specificity and efficiency, with clear identification of TF footprint and signal retention in the targeted cell subpopulations. Furthermore, the deamination reaction showed minimal off-target activity, with high concordance to bulk ChIP-seq reference data. Applied to primary human peripheral blood mononuclear cells (PBMCs), D&D-seq successfully identified CTCF binding sites and enabled integration with advanced machine-learning algorithms for predicting 3D chromatin structure. Furthermore, we integrated D&D-seq with single-cell genotyping to assess the impact of *IDH2* mutations on CTCF binding in a human clonal hematopoiesis sample, uncovering altered binding and chromatin co-accessibility patterns in mutant cells. Altogether, D&D-seq represents an important technological advance enabling the direct mapping of TF or chromatin remodeler binding to the DNA in primary human samples, opening new avenues for understanding chromatin and transcriptional regulation in health and disease.

## INTRODUCTION

Regulatory control orchestrated by chromatin-binding factors, including transcription factors (TFs) and chromatin remodelers, underlies the gene expression programs responsible for maintaining cell identity, executing cellular functions and responding to environmental stimuli. These DNA:Protein interactions are directed through epigenetic features^1^, such as histone modifications and DNA methylation (DNAme), which establish chromatin landscapes, modulating the binding of specific factors, and thereby sculpting the functional genome according to the needs of the cell. Importantly, epigenetic dysregulation has been linked to cellular dysfunction in disease, cancer and aging, where aberrant chromatin landscapes transform the binding landscapes of TFs^2^, altering the normal biological processes of the cell^3^. As such, understanding the complexity and dysregulation of TF binding across different cell types and cell states remains a central challenge in human biology. There are over 1,600 potential TFs encoded in the human genome, with DNA binding motif sequence information, obtained from various *in vitro*, bulk or bioinformatic analyses, established for approximately 1,100 of these^4^. However, DNA sequence motif alone is insufficient for predicting TF binding^4^, which preferentially occurs in open chromatin^5^ and is additionally shaped by TF-TF^6^ as well as TF-nucleosome^7^ interactions. Thus, profiling TF binding in native chromatin contexts is crucial for unraveling the physiologic regulation, and disease-associated dysregulation, of gene expression. However, profiling of these networks across diverse cell types and states is lacking, due to technical limitations of current methods for mapping DNA:Protein interactions in single cells. Chromatin immunoprecipitation with sequencing (ChIP-seq) and cleavage under targets & tagmentation (CUT&Tag) are powerful approaches for mapping genome-wide interactions of target proteins with DNA. These methods identify genomic regions bound by an individual protein of interest through sequencing short DNA fragments that are cross linked or tagmented. In addition, DamID^8,9^ molecular footprinting techniques harnessing DNA methyltransferase activity have been used to map TF binding. However, due to their various protocol requirements, these methods cannot be easily incorporated into available high-throughput single-cell workflows, limiting application to bulk analysis, or in single cells to the profiling of only the chromatin factors that display the strongest interactions with DNA, such as histones. Profiling of TF binding patterns in single cells in primary samples has been mainly restricted to inferential approaches based on expression levels of key downstream TF target genes^10–12^ or through motif analysis of ATAC-seq peaks^13,14^. While analysis of ATAC-seq peaks can be highly informative about the general chromatin landscape, for confident identification of specific TF binding sites, more direct methods are recommended^14,15^. Combining computational TF binding inference analysis with single-cell multi-omic approaches that enable simultaneous capture of chromatin accessibility and gene expression within the same single cell allows for the characterization of genomic regulation across multiple molecular layers (from chromatin landscape to RNA)^16^. Incorporating direct TF binding measurements into such a single-cell framework would provide an unprecedented window into the exact mechanisms driving genome function, transcription networks and pathway regulation. Moreover, this multifaceted analysis spanning different molecular modalities as a readout of direct TF binding would accelerate the identification of key factors that mediate normal cellular activity, as well as those that are disrupted during disease. To address these limitations, we present a new method to accurately map DNA:Protein interactions in low-input samples or single cells that is amenable for direct application to primary human samples. To directly profile TF or chromatin factor binding in single cells, we fused the base-editing deaminase DddA that catalyzes C-to-U single-nucleotide changes to secondary nanobodies targeting an antibody-labeled DNA binding protein of interest, amenable to integration into single-cell genome accessibility workflows such as microfluidic-based scATAC-seq, combinatorial barcoding Paired-seq^17^ or Share-seq^16^, or whole-genome sequencing approaches, such as DLP+^18^ or Primary Template Amplification^19^. As a proof of concept, we incorporate D&D into the 10x Genomics scATAC-seq workflow, enabling the capture of accessible regions of the genome where most gene regulation occurs^5,20^. Upon genome-wide antibody binding to the TF, nanobody-tethered base editor enzyme deaminates proximal cytosines to uracil, leaving a permanent genomic signature at regions of TF binding that can be identified through downstream single-cell sequencing of the tagmented loci. Compatibility with existing droplet-based single-cell sequencing workflows will allow for broad use of Docking and Deamination followed by sequencing (D&D-seq) across systems. We demonstrate the sensitivity and specificity of the D&D enzyme in a deamination assay using DNA oligos or phage DNA, and perform bulk experiments to obtain genome-wide TF binding profiles showing concordance with ChIP-seq data. We test the robustness of single-cell D&D-seq through cell-line mixing experiments, identifying canonical GATA1 binding patterns in erythroid cells. We further apply D&D-seq to primary human peripheral blood mononuclear cells (PBMCs), recovering high enrichment of CTCF binding motifs coinciding with D&D edits. In addition, utilizing single-cell D&D and ATAC data, the D&D-C.Origami pipeline successfully predicts 3D chromatin structure in human primary samples across different cell subtypes. Finally, we integrate single-cell genotyping^21^ into the D&D-seq framework to simultaneously capture genotypes together with TF binding and chromatin accessibility from the same single cell, profiling CTCF binding in an *IDH2*-mutant clonal hematopoiesis of indeterminate potential (CHIP) sample. We envision that the experimental tools and computational analyses developed here can be broadly applied for the interrogation of the interplay between TF or chromatin remodeler binding and chromatin landscapes, as well as defining the impact of somatic mutations on TF binding in single cells, directly from primary samples. We expect that the versatility of D&D-seq will allow for further integration with existing or newly developed single-cell multiomics methods to expand the profiling of downstream modalities including gene expression, opening new avenues for the study of gene regulation in both health and disease.

## RESULTS

### Docking & Deamination (D&D-seq) allows for mapping DNA:Protein binding via molecular foot-printing

CUT&Tag-derived approaches^22,23^, including NTT-seq^24^ and scCUT&Tag-Pro^25^, are suitable for profiling stable and abundant chromatin binders, such as histones, in single cells. These methods are based on CUT&Tag^26^, an approach that generates a library from DNA fragments colocalizing with a chromatin factor of interest by fragmenting out DNA surrounding the antibody-bound chromatin factor. This process is facilitated by pA-Tn5 fusion protein, where protein A (pA) has an affinity to antibodies and Tn5 is a transposase that fragments and tags DNA with sequencing-ready adapters. As Tn5 binds to DNA with high affinity, pA-Tn5 staining and tagmentation are performed under high stringency conditions (e.g., 300-500 mM NaCl) to prevent signal from open chromatin regions mediated by direct binding of Tn5 to DNA. However, these stringent conditions also limit the antibody-tethering activity of the pA-Tn5, resulting in inefficient tagmentation at the chromatin factor-bound sites. In addition, high-salt conditions disrupt weak interactions, such as those between DNA and TFs^27^, or prevent binding of antibodies with low affinity for their targets. As such, these methods are limited in their ability to profile weak DNA:Protein interactions, including those involving TFs and their DNA binding sites. To address this challenge, we developed D&D-seq, a new method for mapping TF binding in single cells. As an alternative approach to tagmentation plus barcoding of fragments at the genomic location bound by the target protein, we reasoned that binding patterns of target proteins can be captured by tethering a base-editing enzyme to a target protein in a precise manner. Briefly, this enzyme is the fusion of the double-stranded DNA deaminase DddA^28,29^ with secondary nanobodies that can recognize species-specific FC domains of immunoglobulins, including rabbit, mouse, goat or rat (referred to as nb-DddA). The fusion protein is thus capable of binding to primary antibodies and catalyzing cytosine deamination in the vicinity of the site that is bound by the targeted TF or chromatin remodeler. This deamination event results in the conversion of cytosine to uracil on the genomic DNA, which can be then identified through sequencing, providing a molecular footprint of the target of interest on the genomic DNA. To showcase the versatility of this approach, we chose to integrate it with the standard ATAC-seq workflow, which provides high-resolution insights into accessible regions of the genome, where most transcription factors bind^5^, making ATAC-seq an ideal method for capturing active genomic regions while simultaneously recording TF molecular footprints (**Fig. 1a)**. To design the fusion enzyme, we selected engineered DddA11 that has increased activity and more versatile DNA editing context compared to the original DddA^29^. To achieve a switch-like control over enzyme activity, we use a split enzyme design, where the deaminase is separated into two polypeptides (**Fig. 1b, Supplementary Fig. 1a, 1b**). This allows for the DddA enzyme to be maintained in an inactive state upon DNA binding, avoiding non-specific deamination due to random interaction of the DddA enzyme with the genomic DNA. The activity of the enzyme can be controlled by the addition of the C-terminal small peptide to the reaction, which through heterodimerization reconstitutes the split enzyme and its catalytic activity^28–30^. We chose the split site with the longest N terminus and linked the nanobodies with the N terminal protein (nb–DddA_NT) so that the C terminal peptide (DddA_CT) is 25 amino acids long (**Fig. 1b**), short enough to be synthesized. We truncated the last 5 amino acids of DddA_CT in the original construct because it is a structurally disordered region (PDB 6u08), and was shown to be functionally redundant by Yin et al^30^. The integration of D&D-seq with ATAC-seq allows profiling of DNA:Protein interactions specifically at regions of open chromatin, where a large fraction of TF binding occurs^5^, while simultaneously providing chromatin accessibility profiling from the same single cells. To first evaluate enzyme base editing specificity, we assessed the deaminase activity of DddA across nucleotide contexts. We tested recombinant nb–DddA_NT activity using a deamination assay. This assay showed that nb–DddA_NT is only active when DddA_ CT is present (**Fig. 1c, Supplementary Fig. 1c, 1d**). Furthermore, we showed that the addition of Zn2+ increases the deaminase activity (**Fig. 1c, Supplementary Fig. 1c, 1d**). Consistent with previous studies^29,30^, we observed that nb–DddA preferentially deaminates cytosines that are in the TpC context (**Fig. 1d**). To assess enzymatic activity, we tested the activity of nb–DddA at different temperatures and incubation times (**Supplementary Fig. 1f, 1g, 1h**), determining that the activity was greater at 37**°**C and 50**°**C compared to 30**°**C and 4**°**C. Together, these results show that our D&D-deaminase recapitulates the base-editing activity of the unmodified enzyme with high fidelity. For bulk D&D-seq, isolated cells are first fixed and permeabilized to allow all the reagents to enter the cell. Then, the sample is incubated with a primary antibody specific to the targeted DNA-binding protein. After washing the unbound antibody, the sample is incubated with nb– DddA_NT. This allows for the deaminase to localize in close proximity with the target. After washing the nb–DddA_NT in excess, we activate the deaminase by incubating the samples with DddA_CT and Zn2+. Followed by washing, the sample can be processed with standard ATAC-seq methods. During the ATAC-seq protocol, genomic DNA is exposed to a highly active transposase (Tn5 or TnY). The transposase simultaneously fragments DNA, preferentially inserts into open chromatin sites, adding the adaptors compatible with downstream amplification and sequencing. Open chromatin is then identified from the sequenced DNA and data analysis can provide insight into gene regulation using the multimodal data. In this way, we profile accessible chromatin and at the same time record the protein binding events on accessible genomic DNA (**Fig. 1a**). To evaluate nb–DddA activity in cells, we next tested nb–DddA in a bulk D&D-seq experiment targeting either CTCF, GATA1 or GATA2 transcription factors in K562 cells. We identified an increased C>T (or G>A) mutation signature in DNA compared to other mutations or to a no-deaminase ATAC control (**Supplementary Fig. 1i, 1j, 1k**). We further characterized the dinucleotide and trinucleotide context in all mutated sites and found that nb–DddA deaminates not only in a TC context, but also in CC and GC contexts (**Supplementary Fig. 1l, 1m**). We reasoned that this is due to high local concentration of DddA tethered around the targeted TFs. To confirm that the deaminase signal is TF-site-specific, we performed motif enrichment analysis using peaks called from reads with DddA-specific edits. Motif enrichment analysis by MEME showed strong CTCF and GATA motifs in the respective peaks (**Fig. 1e, 1f, 1g**). A deamination footprint analysis of edit events in a 200 base-pair window around all the HOCOMOCO-defined^31^ CTCF, GATA1 or GATA2 binding sites in the genome showed the expected bimodal distribution of C edit events, where the binding sites for the targeted TF are at the center of the two modes, with low editing frequency in background accessible regions lacking binding sites (**Fig. 1h, 1i, 1j**). Furthermore, we used ENCODE^20^ CTCF or GATA1 ChIP-seq data from K562 cells to define on-target and off-target peaks, and identified significantly higher deaminase activity in on-target peaks compared to the off-target peaks or to the background (**Fig. 1k**). These results support the efficient activity of nb–DddA and provide evidence that TF binding can be profiled with high specificity in native chromatin conditions.

**Fig. 1.**
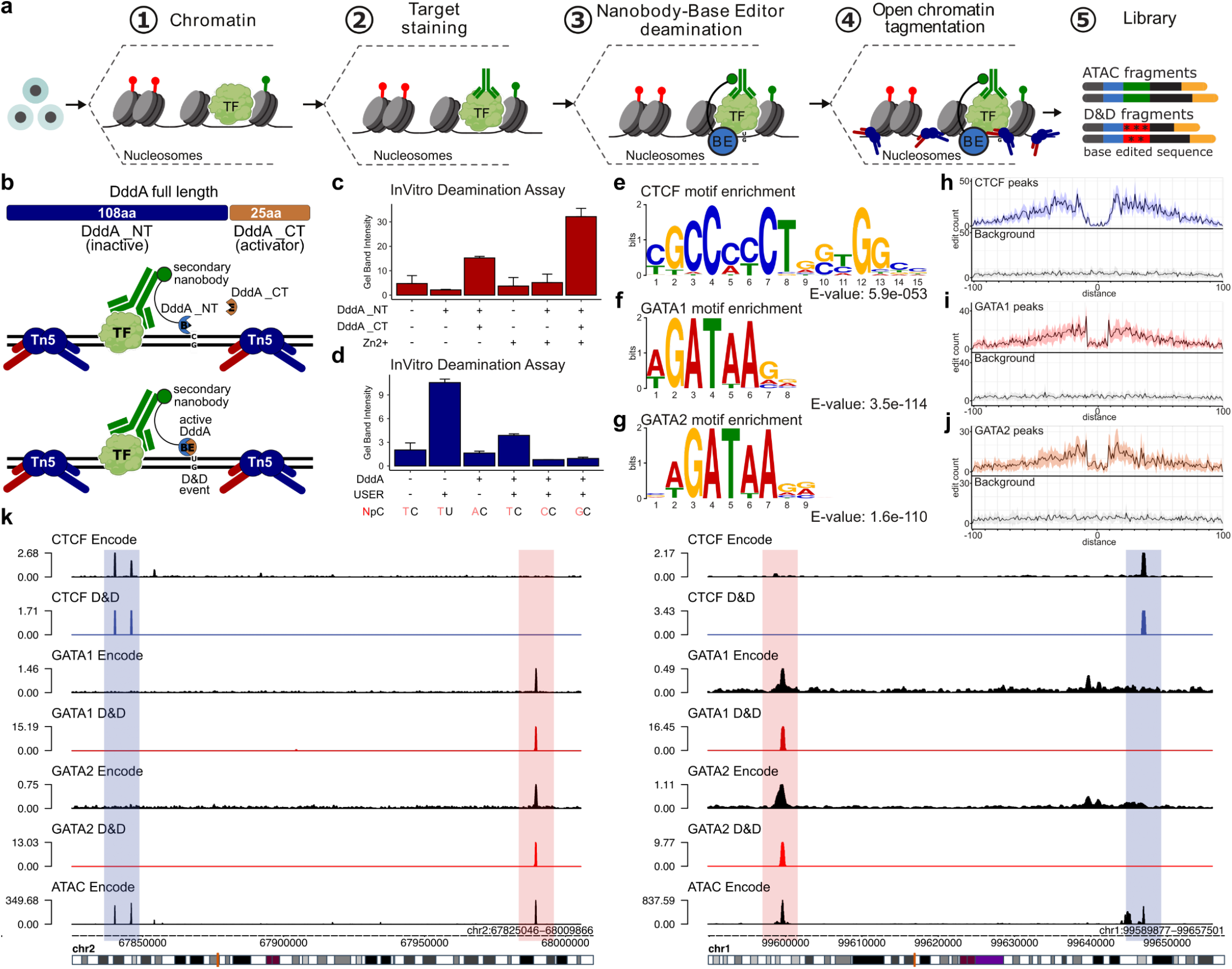
Docking & Deamination allow for transcription factors binding map via molecular foot-printing. **a)** Schematic illustration of D&D-seq technology. TF: transcription factor. BE: base editor (deaminase). **b)** Illustration of the split DddA base editor comprising N-terminal (DddA_NT) and C-terminal (DddA_CT) regions. DddA_NT is fused to nanobodies and is activated by addition of the C-terminal peptide. **c)** In vitro deamination assay (Supp. Fig. 1c) to quantify deamination efficiency in the presence or absence of Zn2+. The bars represent the average DNA intensity (n = 2) of the cleaved product quantified by ImageJ for split DddA11 Rb-DddA_NT, DddA_CT, or both. **d)** In vitro deamination assay (Supp. Fig. 1c) to quantify deamination efficiency for the split DddA11 deaminase and the uracil-targeting USER enzyme in different dinucleotide contexts. The bars represent the average DNA intensity (n = 2) of the cleaved product quantified by ImageJ. **e)** Motif enrichment analysis of bulk D&D-seq for CTCF in K562 cells. Genome-wide D&D-seq reads were analyzed for enrichment of the CTCF binding motif using MEME. **f)** Same as (e) for GATA2. **g)** Same as (e) for GATA2. **h)** Molecular footprint of bulk D&D-seq for CTCF in K562 cells showing count of C-to-U edits at aggregated CTCF sites compared to a background ATAC region with no CTCF binding site. **i)** Same as (h) for GATA1. **j)** Same as (h) for GATA2. **k)** Genome browser tracks for representative regions of the human genome (left, chr 2 region; right, chr 1 region). D&D-seq was performed on K562 cells for CTCF (light blue), GATA1 or GATA2 (red). Encode ChIP-seq data for the 3 proteins (black) are used as reference. Sequencing data were normalized as bins per million (BPM) mapped reads.

### D&D-seq allows for single-cell DNA:Protein binding profiling

We next tested the ability of D&D-seq to record TF binding in single cells when incorporated into the 10x Genomics single-cell (sc) ATAC-seq workflow. We performed cell line mixing experiments, selecting two distinct cell lines (lymphocyte cell line CA46 and erythroblast cell line K562) and two TF targets (CTCF and GATA1) to allow us to assess the targeting specificity and potential cross-contamination of signal in D&D-seq. The two cell lines were individually crosslinked, permeabilized, and stained with a CTCF antibody for CA46 and a GATA1 antibody for K562. Cells were then washed, equally mixed and incubated in the D&D-seq buffer containing the activating C-terminal peptide of DddA and the Zn2+ cofactor necessary for the activation of the deamination reaction. In this manner, we are able to record the presence of CTCF or GATA1 on the genomic DNA of each of the tested cell lines. After this step, the sample can be processed with custom or commercially available single-cell ATAC workflows with minimal modifications to the original protocol. We profiled K562 (n = 1,732 cells) and CA46 (n = 4,108 cells) (**Fig. 2a**), obtaining 5,263 +/− 2,633 (mean +/− standard deviation) fragments per cell for K562 and 5,437 +/− 2,588 (mean +/− standard deviation) fragments per cell for CA46 (**Fig. 2b**), reflecting high quality tagmentation data (**Supplementary Fig. 2a, 2b, 2c**). Cell lines were identified by gene accessibility score of known marker genes (**Fig. 2c**). We projected cells into a low-dimensional space using latent semantic indexing (LSI) and uniform manifold approximation and projection (UMAP) using Seurat, and clustered cells using a shared nearest neighbor (SNN) approach. There were two clear clusters corresponding to the two cell types showing good separation between CA46 and K562 cells, as defined by highly accessible markers for lymphoid (**Fig. 2d**) and erythroid cells (**Fig. 2e**), demonstrating that the D&D-seq reaction does not affect the overall performance of the scATAC. Importantly, at the single-cell level, we were able to map protein binding events in all the cells analyzed, suggesting that the deamination reaction has a relatively high efficiency (**Methods**). While the absolute number of edits per cell is relatively low (**Fig. 2f, 2g, 2h, 2i**), this is comparable to established tagmentation-based single-cell transcription factor binding inference methods. Furthermore, the data can be effectively integrated through pseudobulking or meta-cell analysis, enabling robust downstream interpretations. Since the antibody staining step was performed before mixing the two cell lines, we were able to assess the presence of cross-contamination potentially occurring during droplet encapsulation, barcoding, and library preparation reactions. Our results showed that the signals for CTCF and GATA1 are mutually exclusive and are retained exclusively in the subpopulation stained with the respective specific antibody, demonstrating that the specificity of the signal is maintained at the cellular level (**Fig. 2f, 2g, 2h, 2i**). To provide support for the specificity of D&D-seq, we performed de novo motif discovery analysis on the pseudo-bulk D&D-seq mutational peaks (**Fig. 2j, 2k, 2l, 2m**). We observed significant enrichment of the CTCF binding site (e-value 1.3e-171) in CA46 cells (**Fig. 2j**) and the GATA1 binding site (e-value 2.1e-89) in K562 cells (**Fig. 2l**). We next orthogonally validated our findings using a motif-centric, rather than de novo, approach. Here, we used the total reads obtained from the single-cell experiment to calculate the frequency of D&D events in a 200 base-pair window around all the HOCOMOCO-defined^31^ CTCF or GATA binding sites in the genome, and compared this with the frequency of C-to-T events observed in all the accessible peaks measured in our experiment. This analysis revealed the presence of the expected bimodal distribution of C deamination events, where the binding sites for the protein of interest represent the center of the two modes (**Fig. 2k, 2m**, top). However, when the same analysis was performed for accessible regions lacking known binding sites, we observed negligible edit event frequency and the absence of clearly-defined distribution patterns (**Fig. 2k, 2m**, bottom). Moreover, when we repeated this analysis but for the non-targeted factor for each cell type, we observed no GATA1 binding signal in CA46 cells and no CTCF binding signal in K562 cells (**Supplementary Fig. 2d**). These results demonstrate across two TFs that our D&D-seq approach faithfully recapitulates TF binding at the single-cell level. We next sought to evaluate the specificity of the deamination reaction at the site of target protein binding. To do so, we collapsed the reads from a specific cluster to obtain pseudo-bulk alignments. We next extracted the reads where we identified C-to-T conversion events and generated a D&D coverage plot. The peaks obtained with our *in situ* footprinting approach showed high concordance with bulk ChIP-seq reference data obtained from the ENCODE consortium^20^ (**Fig. 2n, 2o**), suggesting that base editing events occur only in the proximity of the targets of interest, with minimal off-target activity. We quantified the similarity between our single-cell D&D-seq reads and those from ENCODE, showing high correlation between CTCF signals from each data type and between GATA1 signals for each data type, but low correlation across the CTCF and GATA1 signals (**Fig. 2o**). We note that the correlation values we observed between our single-cell D&D-seq reads and those from ENCODE were comparable to the results obtained using well-established technologies on histones, such as CUT&Tag on H3K27me3 or H3K4me3^26^, which is often used as a benchmarking target due to its high abundance and high-quality results (**Supplementary Fig. 2e**). The lack of technologies with similar capabilities, especially for single cell applications, limits direct benchmarking of D&D-seq with the current state-of-the-art methods for profiling DNA: Protein interactions. Despite this gap, attempts have been made to profile the chromatin occupancy of non-histone proteins in low input samples by modifying well-established bulk protocols. Among those, ultra-low input cleavage under target and release using nuclease (uliCUT&RUN) is a variant of CUT&RUN, with key modifications to reduce background signal, increase output and decrease the amount of starting material required to generate protein occupancy profiles from mammalian cells^32^. This protocol was used to profile the genomic locations of the insulator protein CTCF from populations of mouse embryonic stem cells (mESCs) ranging in number from 500,000 to 10^33^. In order to provide further benchmarking for D&D-seq, we iteratively randomly subsampled (n=100) the single-cell CA46 subpopulation, stained with the CTCF antibody, to generate matching datasets with the same number of cells analyzed by uliCUT&RUN. As negative control, the same number of K562 cells, which represent cells that were not stained with CTCF antibody, were subsampled to match the uliCUT&RUN negative control, cells that were not stained with antibody. To evaluate the specificity of the two protocols, we calculated the fraction of the reads in the peaks (FRiP), using a reference peak annotation obtained from a high-quality ChIP-seq. A high fraction of reads in peaks indicates that the majority of the reads are located in the regions of interest, and that the experiment has a high signal-to-noise ratio. On the other hand, a low fraction of reads in peaks may indicate that the majority of the reads are located in non-specific regions, and that the experiment has a low specificity. D&D-seq data showed that the majority of the base-edited reads map to regions where CTCF was previously mapped, consistently reaching ∼60% FRiP across analyses. Moreover, the signal-to-noise ratio was largely unaffected by the number of the cells analyzed, showing similar values for the 5,000-, 500-, 50-, and 10-cell sample sizes, with mean FRiP equal to 56.75%, 56.78%, 56.83%, and 57.00% respectively (**Fig. 2p**, top). As expected, the cell line that was not stained with the CTCF antibody showed very low values of FRiP in all the input conditions, indicating that the majority of the edited reads were not specific for CTCF. Importantly, at least for this specific metric, D&D-seq outperforms uliCUT&RUN, which scored 13.69%, 13.69%, 13.89%, and 14.04% FRiP for the 5000-, 500-, 50-, and 10-cell sample sizes, reflecting a majority of reads mapped to regions of the genome that are not bound by CTCF (**Fig. 2p**, bottom). In conclusion, these data confirm the specificity of the scD&D-seq signal, indicating that our method performs at a level comparable to the current state-of-the-art approaches. However, beyond achieving similar accuracy, D&D-seq offers significant advantages, as it is capable of simultaneously capturing a wider range of biological features at single-cell resolution, thus enabling a deeper exploration of complex regulatory mechanisms that were previously inaccessible.

**Fig. 2.**
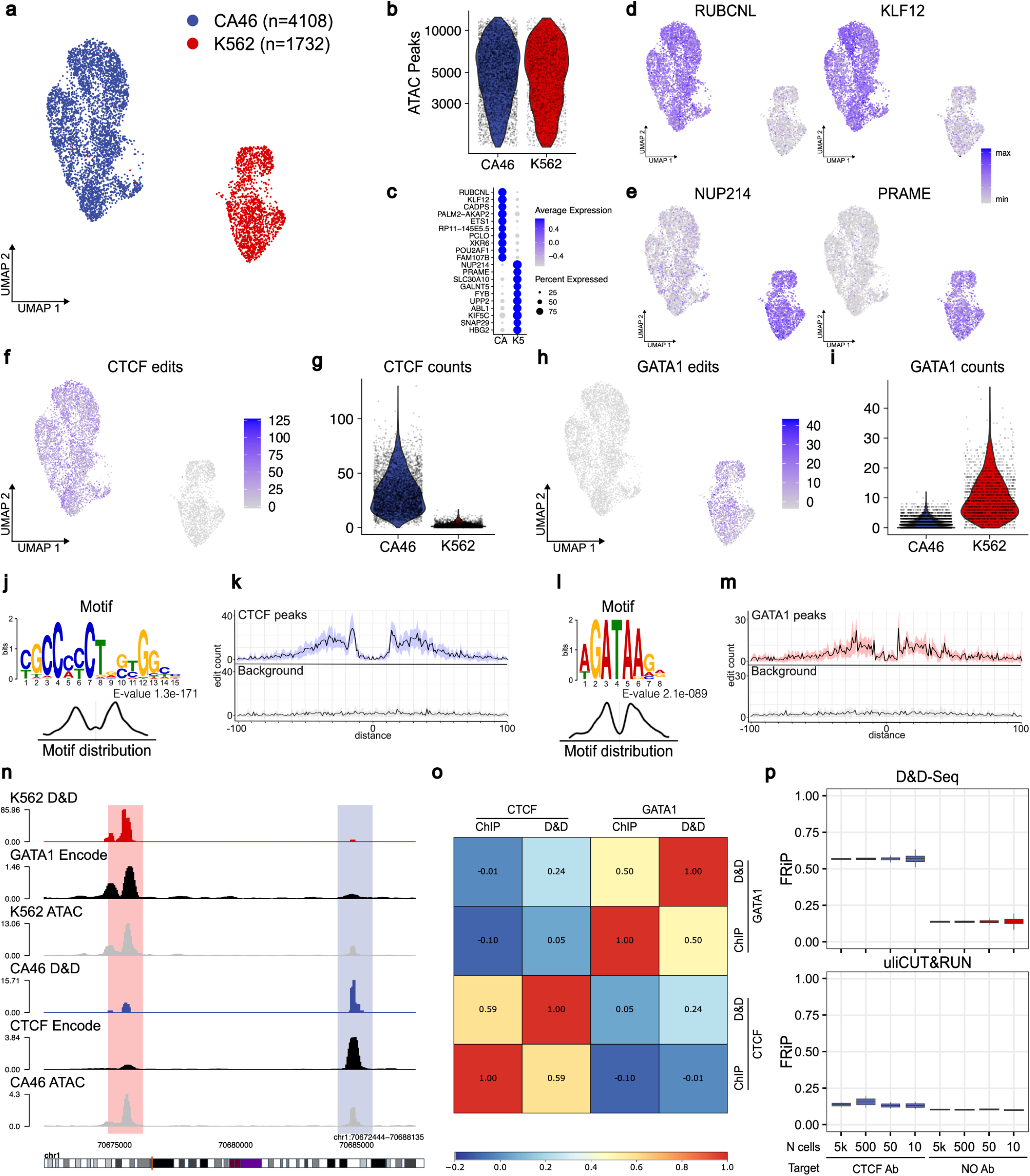
D&D-seq allow for single cell mapping of DNA:Protein interaction. **a)** UMAP of cell line mixing study of CA46 (n = 4,108 cells) and K562 (n = 1,732 cells) cells in the ATAC space. **b)** Violin plot showing the number of ATAC fragments per cell for each subpopulation from (a). **c)** Accessibility scores for marker genes defining the subpopulation lineage (FDR < 0.05). **d)** UMAP projection of gene accessibility score for CA46 marker genes *RUBCNL* and *KLF12*.**e)** UMAP projection of gene accessibility score for K562 marker genes *NUP214* and *PRAME*.**f)** Projection of CTCF D&D edit events into the UMAP space. **g)** Violin plot of CTCF D&D edit events in CA46 and K562 cells. **h)** Projection of GATA1 D&D edit events into the UMAP space. **i)** Violin plot of GATA1 D&D edit events in CA46 and K562 cells. **j)** Motif enrichment analysis for D&D-seq reads in CA46 cells. **k)** Molecular footprint of pseudo bulk D&D-seq for CTCF in CA46 cells showing count of C-to-T edits at aggregated CTCF sites compared to a background ATAC region with no CTCF binding site. **l)** Motif enrichment analysis for D&D-seq reads in K562 cells. **m)** Molecular footprint of pseudobulk D&D-seq for GATA1 in K562 cells showing count of C-to-T edits at aggregated GATA1 sites compared to a background ATAC region with no GATA1 binding site. **n)** Pseudobulk genome browser tracks for representative regions of the human genome (chr1). scD&D-seq was performed on a mixture of K562 and CA46 cells for GATA1 (red) or CTCF (light blue), respectively. Encode ChIP-seq data for the 2 proteins (black) as reference. ATAC peaks are in gray. Sequencing data were normalized as bins per million (BPM) mapped reads. **o)** Genome-wide Pearson correlation between pseudobulk scD&D-seq and Encode ChIP-seq for CTCF and GATA1 in K562 cells. Peaks are defined by ENCODE ChIP-seq for CTCF and GATA1. **p)** Fraction of fragments falling in ENCODE ChIP-seq peak regions for CTCF for D&D-seq (top) and uliCUT&RUN (bottom). No antibody (Ab) control indicates cells that were not stained with the CTCF antibody.

### Single-cell DNA:Protein interaction profiling in primary human cells

We tested D&D-seq for analysis of primary human PBMCs. We selected CTCF as an initial target due to its ubiquitous presence on the genome in all cell types and cell states, and the presence of a defined consensus DNA binding sequence^34^. For this experiment, mobilized PBMCs collected from healthy donors were crosslinked, permeabilized, and stained with a CTCF-specific polyclonal antibody. The sample was then processed with the D&D reaction, followed by standard 10x Genomics scATAC with minimal modification (**see Methods**). We obtained high-quality single cells (n = 5,358 cells), with 15,534 +/− 5,934 (mean +/− standard deviation) fragments per cell. We projected cells into a low-dimensional space using LSI and UMAP, clustering cells using the chromatin accessibility data (**Fig. 3a**). We assigned cell states based on chromatin accessibility profiles of established marker genes and retrieved all the expected subpopulations, demonstrating that the addition of the D&D reaction does not impact the overall quality of the chromatin accessibility assay (**Supplementary Fig. 3a, 3b**). Critically, we mapped CTCF binding sites in this complex primary tissue (**Fig. 3b, Supplementary Fig. 3c**). We extracted the reads with in situ C-to-T transition labeling and identified CTCF binding sites in close proximity to C-to-T deamination events (**Fig. 3c, 3d**). These results demonstrate the direct applicability of D&D-seq in primary human samples. We next speculated that the availability of chromatin accessibility and CTCF binding from the same single cells could be used to enhance machine-learning based algorithms developed to predict chromatin 3D structure in bulk, paving the way for these methods to be used by the single-cell genomics field. In particular, we focused on the C.Origami pipeline^35^, which utilizes chromatin accessibility data coupled with CTCF ChIP-seq data to feed a deep neural network that performs de novo prediction of cell-type specific 3D chromatin organization. We took advantage of our D&D-seq protocol to generate a cell-type specific CTCF binding profile that can be used as a genomic feature by the C.Origami algorithm to generate a single-cell pseudo-bulk Hi-C genomic contact map. The D&D-C.Origami pipeline successfully generated a contact matrix by using scD&D and scATAC data as input. The predicted structures closely recapitulate publicly available experimental Hi-C data. Both short and long-range interactions were efficiently predicted by D&D-C.Origami in human primary CD8 T cells (**Fig. 3e**). To validate and quantify the accuracy of the D&D-C.Origami approach, we generated 3D chromatin predictions for 100 random 2-Mb chromatin regions in all major cell subtypes (including B cells and monocytes, **Supplementary Fig. 3d, 3e, 3f, 3g**). The input data consisted of ATAC-seq regions and either CTCF ChIP-seq or CTCF D&D-seq. 3D chromatin inference without CTCF information served as a negative control. The C.origami pipeline was able to infer 3D chromatin structure with high accuracy, exhibiting a high correlation with experimental data, with comparable accuracy using scD&D-seq data instead of ChIP-seq data (**Fig. 3f, Supplementary Fig. 3e, 3g**). We thus envision that scD&D-seq will enable the broad exploration of chromatin 3D structure changes during major remodeling events, such as changes in differentiation state or aging.

**Fig. 3.**
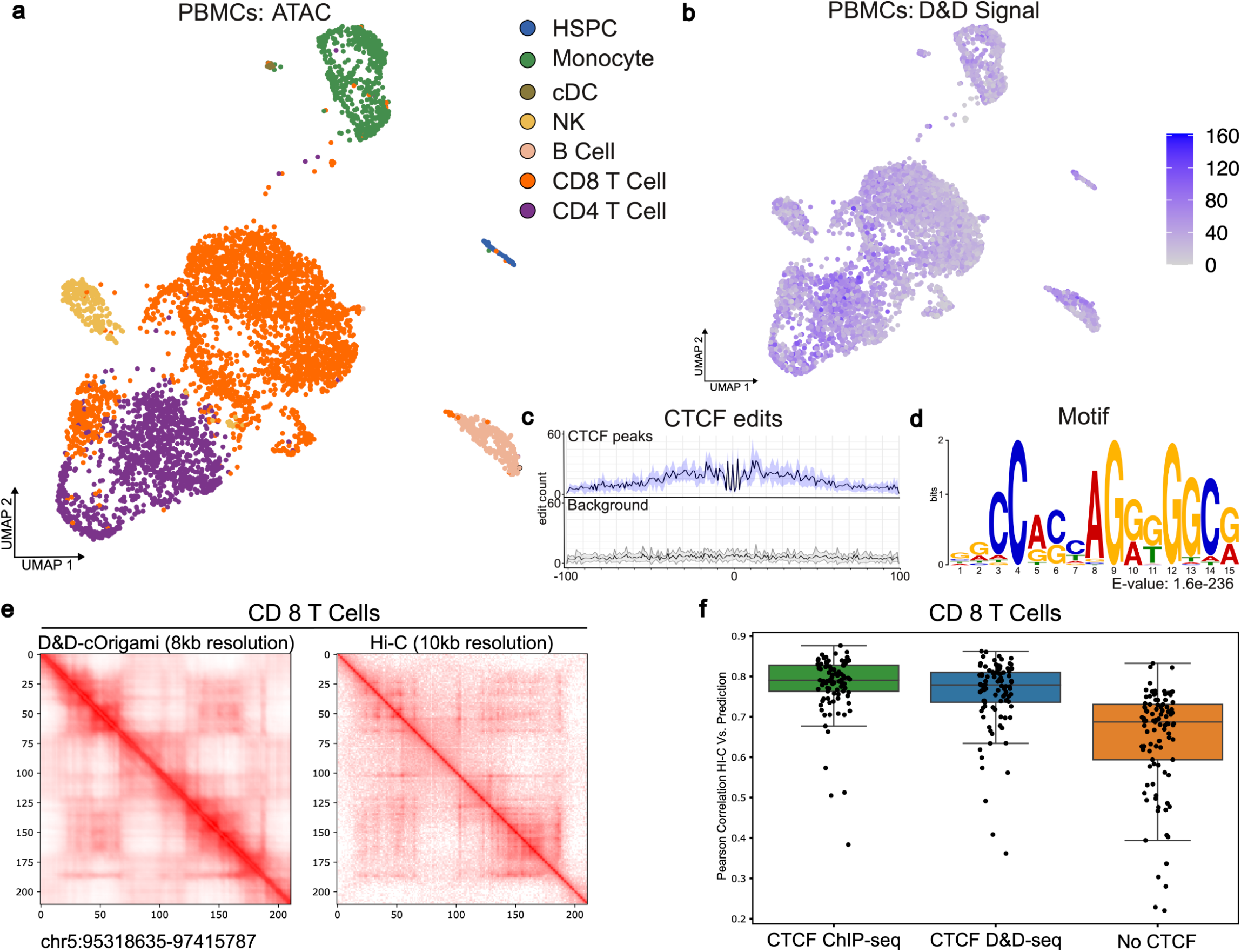
D&D-seq allows for DNA:Protein interaction mapping and 3D chromatin inference in primary human cells. **a)** UMAP of primary PBMCs study (n = 5,358 cells) in the ATAC space. HSPC, hematopoietic stem/progenitor cells; cDC, conventional dendritic cells; NK, natural killer cells. **b)** Projection of CTCF D&D edit events into the UMAP space. **c)** Distribution of C to U D&D events around CTCF binding site (top) and ATAC (bottom) across all the genome. **d)** Motif enrichment analysis for D&D-seq reads. **e)** D&D-C.Origami-predicted (left) and experimental (right) Hi-C matrices of CD8 T cells. **f)** Pearson correlation of 100 randomly chosen CD8 T cells regions experimental Hi-C data (ENCODE) and: C.Origami 3D chromatin predictions generated with CTCF ChIP-seq data (ENCODE), CTCF D&D-seq or in the absence of CTCF information. Boxes represent the interquartile range, lines represent the mean, and error bars represent the 1.5 interquartile ranges of the lower and upper quartile.

### Linking genotype to DNA:Protein interaction profiles and chromatin landscapes in single cells

Given the compatibility of the D&D workflow with high-throughput single-cell platforms such as 10x Genomics, it is possible to incorporate this versatile backbone technology into other single-cell multiomics frameworks that capture multiple modalities. To this end, we reasoned that integrating single-cell genotyping capability would enable the extension of D&D-seq to study the effects of somatic variants on gene regulation. Given the mixture of mutated and wild-type cells within clonally mosaic tissues, single-cell multi-modal methods that capture both genotype and phenotype together are required to provide biological insights into mechanisms of clonal outgrowth in normal, aging or diseased tissues^36^. To address this challenge, we previously developed Genotyping of Transcriptomes (GoT)^37^ and Genotyping of Targeted Loci and Chromatin Accessibility (GoT-ChA)^21^ to enable high-throughput single-cell genotyping together with transcriptome profiling or chromatin accessibility profiles. Leveraging this conceptual framework for linking single-cell genotype to molecular phenotype, we incorporated our D&D-seq approach into single-cell genotype-aware multi-omics, to link somatic mutations to changes in TF binding, and applied this workflow to profile a human CHIP sample. Briefly, in GoT-ChA, isolated cells are subjected to transposition of genomic DNA (gDNA) and loaded into microfluidics devices (**Supplementary Fig. 4a, steps 1-4**). During cell barcoding reactions, additional gene-specific primers are added to capture a locus of interest with an in-droplet PCR reaction, with a handle that is compatible with the 10x scATAC platform (**Supplementary Fig. 4a, step 5**). The product is then split, with an aliquot used for an amplicon genotype library and the remaining 90% used for scATAC library construction and sequencing (**Supplementary Fig. 4a, steps 6-7**). The scATAC and genotyping libraries can then be analytically integrated via shared cell barcodes, linking chromatin accessibility to targeted genotyping at single-cell resolution. To incorporate genotyping into the D&D-seq framework, GoT-ChA primers are added during the scATAC in-drop PCR step. To test D&D-GoT-ChA, we profiled CTCF binding in PBMCs from a patient with CHIP carrying an *IDH2*^*R140Q*^ mutation. The mutation was identified by targeted panel sequencing and had a variant allele frequency of 0.15. Integration of GoT-ChA with D&D-seq allowed us to build a multiomic dataset that includes the genotype of individual cells, along with accessible chromatin status and CTCF binding pattern. The simultaneous implementation of targeted genotyping and D&D-deamination did not affect the quality of the chromatin accessibility data, which we used to resolve the tissue into highly granular cell populations, identifying all the major cell subtypes expected for this sample (**Fig. 4a**). The experiment was performed in two technical replicates, and after quality check and batch correction, the data from the individual experiments were merged into a single dataset of 15,807 high-quality cells. The resulting dataset showed appropriate QC metrics, with a mean of 7,592 fragments per cell (+/− 3,971, standard deviation), enrichment around transcriptional start sites, and expected library size distribution (**Supplementary Fig. 4b**). Accessibility in proximity of key marker genes was used to guide cell annotation, allowing us to identify all the major cell types (**Supplementary Fig. 4c**). The genotyping library obtained by GoT-ChA targeted amplification was sequenced and processed following our previously published pipeline (https://github.com/landau-lab/Gotcha)^21^. We successfully genotyped 33.59% of single cells (5,291 out of 15,807 cells), identifying *IDH2*^*R140Q*^-mutant and wild-type cells. Genotyping data were assigned to individual cells and projected into the accessibility latent space to visualize the distribution of wild-type and mutant cells in the tissue (**Fig. 4b**). This analysis revealed an enrichment of mutant cells in the CD8 T cell subcluster, showing that the *IDH2* mutation was almost exclusively found in this specific cell type (**Fig. 4c, Supplementary Fig. 4d**). To validate this finding, peripheral blood cells from the same donor were immunolabeled and split into the main subpopulation by FACS (**Methods, Supplementary Note**). Genomic DNA was isolated from natural killer (NK), monocytes, CD8 T cells, CD4 T cells, and B cells and processed for standard genotyping by bulk nanopore sequencing. The bulk genotyping data matched the single-cell genotyping data obtained with D&D-GoTChA (**Fig. 4d**), further validating the precision of our single-cell approach. Next, we analyzed the CTCF binding pattern across cells. The CTCF molecular footprinting signal was present in all the single-cell clusters and evenly distributed across all the cell types, as expected from this factor (**Fig. 4e, 4f**). We identified 5,631 CTCF positive peaks, and the motif enrichment analysis of the footprinted reads again revealed the specificity of our deamination assay, showing a significant enrichment of CTCF binding sites in the proximity of editing events (**Fig. 4g, Supplementary Fig. 4e**). IDH-mutant cells have high levels of 2-hydroxyglutarate (2-HG) that interfere with the TET family of 5′-methylcytosine hydroxylases^38^. TET enzymes catalyze a key step in the removal of DNA methylation^39^. Previous in vitro studies have shown that IDH-mutant cells exhibit hyper-methylation at CTCF binding sites, compromising the binding of this methylation-sensitive insulator protein^40,41^. Reduced CTCF binding is associated with loss of insulation between topological domains and aberrant gene activation^40^, although the extent and functional significance of IDH-mutant mediated alterations of epigenetic states remain unclear in vivo. Our D&D technology allows us, for the first time, to address this question by analyzing differences in CTCF binding between IDH wild-type and mutant cells in primary human cells. As the vast majority of mutant cells were detected in CD8 T cells, we focused our analyses on this specific cell type (**Fig. 4h**). First, we observed that CD8 T cells form two distinct clusters characterized by the differential presence of wild-type and mutant cells. Therefore, we reassigned the identity of these two clusters as CD8 wild-type enriched and CD8 mutant enriched clusters to assess *IDH2*-driven phenotypic changes **(Fig. 4h, Supplementary Fig. 4f)**. We assessed CTCF binding in CD8 T cells by comparing the CTCF D&D binding signals (C>T edit read counts) between *IDH2* wild type (WT) and mutant (IDH2^R140Q^, MUT). Notably, the vast majority of statistically significant binding sites differentially bound by CTCF exhibit a decrease in the CTCF binding signal between mutant and wild-type cells (**Fig. 4i**), which corroborates previous reports suggesting that CTCF binding is reduced in *IDH*-mutant glioma and acute myeloid leukemia, mediated by DNA hypermethylation at CTCF binding sites^40,41^. For example, the binding site in close proximity to the *GIT1* gene showed the strongest reduction in binding. GPCR-kinase interacting protein 1 (GIT1) is a scaffold protein that interacts with proteins such as RAC1, PAK, and paxillin to regulate actin cytoskeleton remodeling^42^. This is crucial for T cell migration, synapse formation, and activation^43,44^. Furthermore, this protein has been implicated in modulating T cell-mediated inflammatory responses^45^. To further explore the functional consequences of CTCF loss in this genomic domain, we took advantage of our ability to simultaneously profile and integrate differential CTCF binding, chromatin accessibility, and genotyping to identify changes in cis-regulatory DNA interactions affecting the *GIT1* topologically associated domain (TAD) that may be a consequence of *IDH2* mutation. To evaluate the potential alterations in 3D genomic interactions between mutant and wild-type cells, we once again utilized the C.Origami algorithm^35^ to generate single-cell pseudo-bulk Hi-C genomic contact maps based on our CTCF D&D-seq profiles. Notably, we observed a significant alteration of 3D genome contacts in mutant-enriched T cells compared to wild-type-enriched T cells at the *GIT1* locus. In greater detail, wild-type cells tended to form structures resembling canonical TADs (**Fig. 4j**, red pixels), while the interactions formed by mutant cells were more evenly distributed throughout the entire genomic region (**Fig. 4j**, blue pixels). To evaluate how aberrant CTCF binding in this region may affect gene regulation, we utilized the Cicero algorithm^46^ to evaluate the co-accessibility of regulatory elements present in the chromatin hubs in proximity to the perturbed CTCF elements. Here, we expanded the genomic window from the *GIT1* locus to include additional up-/downstream CTCF binding sites that were bound by CTCF in both wild-type and mutant cells. A comparison of co-accessible peaks between wild-type and mutant T cells in this region, showed that the chromatin architecture was substantially reshaped in mutant T cells (**Fig. 4k, 4l**) across four main accessible regions (red boxes). To further visualize this observation, we compared the pairwise co-accessibility landscape of the *GIT1* TAD by plotting smoothed co-accessibility scores of all the possible pairwise interactions of ATAC peaks in this region. In wildtype cells these interactions appeared more well-defined, while in mutant cells that lost CTCF binding events around the GIT1 locus, co-accessibility patterns appeared more disorganized (**Fig. 4m, 4n**). The co-accessibility periodicity observed in wild-type cells appears compromised in mutant cells, likely reflecting a less defined topological domain organization, perhaps due to the loss of CTCF binding and its insulator function, which may prevent active genomic loci from interacting in a more random manner (**Fig. 4n**). Similarly, we observed the same behavior at the *CNIH2* locus, where well-defined co-accessibility in wild-type cells became disorganized in mutant cells due to the loss of CTCF binding (**Supplementary Fig. 4h, 4i, 4j, 4k**). This consistent pattern implies that the mutation-driven disruption of chromatin organization is not limited to a specific locus but rather reflects a broader phenotype resulting from the presence of IDH2 mutation, and consequently disruption of CTCF binding. These observations align with *in vitro* studies performed by acute depletion of CTCF, which demonstrated that CTCF is essential for looping between CTCF target sites and the insulation of TADs^47^. Moreover, additional interactions were observed with an upstream peak within *NUFIP2* gene locus in the mutant T cells, while absent in the wild type (**Fig. 4k-n**). Thus, in wildtype cells, where both CTCF binding sites in the region are bound by CTCF protein, we observe a more coordinated orchestration of co-accessibility within small genomic area and limited interactions with upstream regions (**Fig. 4m**). In mutant cells, however, co-accessible peaks in the *GIT1* TAD were disrupted with loss of the CTCF insulator function and formed new interactions with upstream of *GIT1*, especially with *NUFIP2* gene, suggesting the formation of a new gene regulatory network upon loss of CTCF binding in this locus (**Fig. 4l**). NUFIP2 was previously reported to regulate *ICOS* expression in mice, the inducible T-cell co-stimulatory receptor which is a critical regulator of T cell proliferation and cytokine production.

**Fig. 4.**
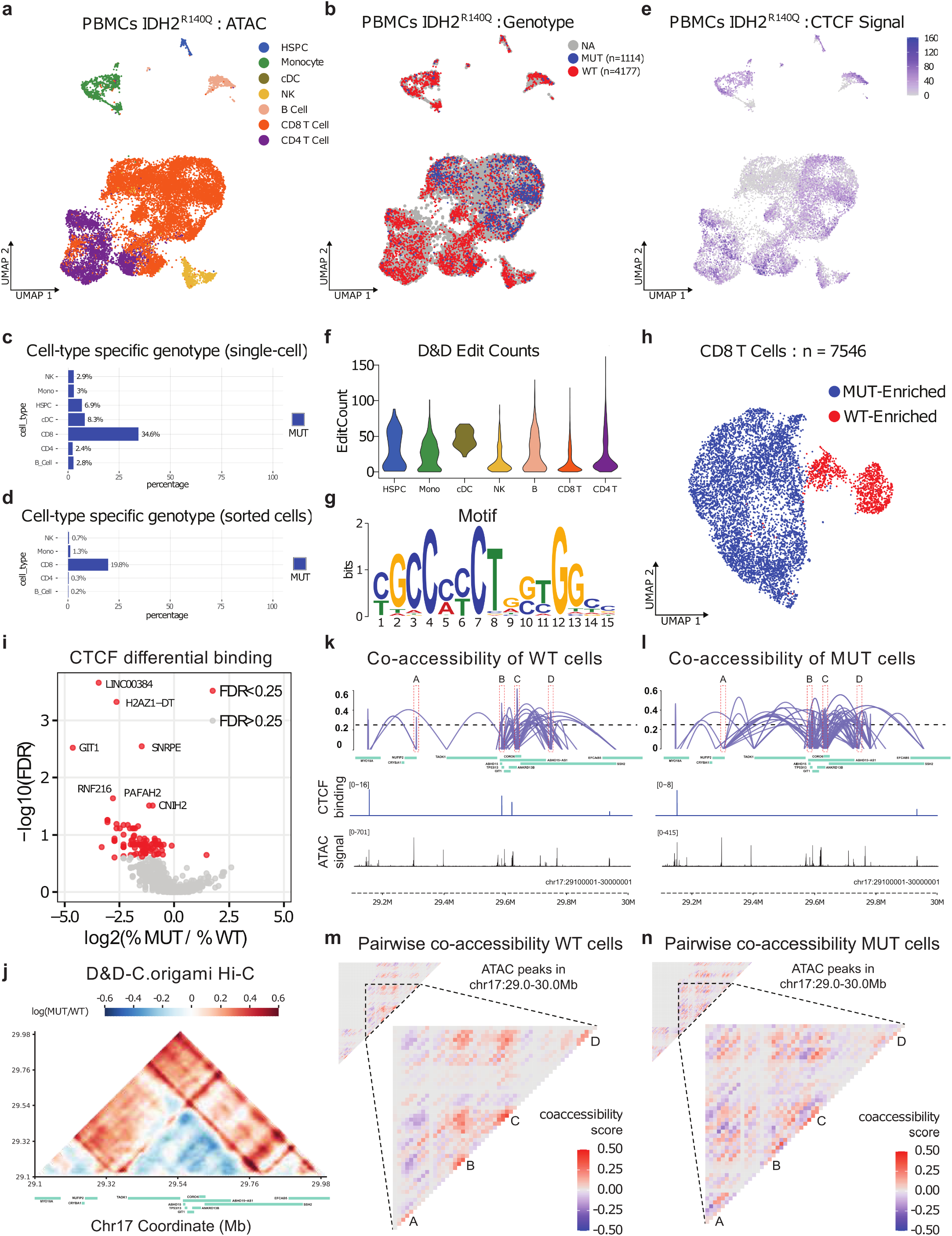
Multiomic application of D&D-seq to human primary clonal hematopoiesis samples. **a)** Integrated chromatin accessibility uniform manifold approximation and projection (UMAP) after reciprocal latent semantic indexing (LSI) integration of patient samples (n = 2 replicates, 15,807 cells) illustrating the expected hematopoietic populations. **b)** Integrated UMAP colored by GoT-ChA-assigned *IDH2* genotype as wild type (WT; n = 4,177 cells; blue) or mutant (MUT; n = 1,114 cells; red) and not assignable (NA; n = 10,516 cells; grey). **c)** Percentage of cells genotyped as *IDH2* mutant (MUT; red) for each cell subtype. **d)** Validation of GoT-ChA results showing percentage of mutant cells genotyped using bulk targeted sequencing for *IDH2* in each cell subtype.**e)** Projection of CTCF D&D edit events into the UMAP space. **f)** Number of D&D edit events recorded for each cell subtype. **g)** Motif enrichment analysis for D&D-seq reads. **h)** Uniform manifold approximation and projection (UMAP) of the CD8 T cell fraction of the sample (n = 7,546 cells) with genotypes annotated as mutant-enriched or wild type-enriched. **i)** Differential CTCF binding regions in mutant versus wild-type CD8 T cells. Total of 1,102 CTCF peaks with more than five D&D edits and five cells were tested using LMM (see Methods). Gene regions with false discovery rate < 0.25 are highlighted in red. **j)** Heatmap displaying D&D C.Origami Hi-C predicted interactions enriched in mutant versus wild-type CD8 T cells at the genomic region on chr 17 encompassing the *GIT1* locus. **k)** Co-accessible regions determined from ATAC data (purple lines) in wild-type CD8 T cells at the *GIT1* locus. CTCF binding events and ATAC signal detected with D&D-seq are displayed in the bottom tracks. **l)** Co-accessible regions determined from ATAC data (purple lines) in mutant CD8 T cells at the *GIT1* locus. CTCF binding events and ATAC signal detected with D&D-seq are displayed in the bottom tracks. **m)** Heatmap displaying pairwise ATAC-seq peak co-accessibility in wild-type CD8 T cells at the region corresponding to k and l co-accessibility plots (top-left). Zoom-in heatmap displaying pairwise ATAC-seq peak co-accessibility within a, b, c, and d peaks (bottom-right). Co-accessibility score was smoothed by averaging three adjacent peaks. **n)** Heatmap displaying pairwise ATAC-seq peak co-accessibility in mutant CD8 T cells at the region corresponding to k and l co-accessibility plots (top-left). Zoom-in heatmap displaying pairwise ATAC-seq peak co-accessibility within a, b, c, and d peaks (bottom-right). Co-accessibility score was smoothed by averaging three adjacent peaks.

## DISCUSSION

Single-cell mapping of chromatin factors has the potential to uncover the rules governing gene network regulation directly within individual cells, broadening our ability to understand cell biology and identify specific vulnerabilities within disease states. However, existing single-cell compatible chromatin profiling technologies, such as NTT-seq^24^ and scCUT&Tag^26^, require complex workflows and very stringent buffer conditions to avoid non-specific tagmentation that could result in unusable datasets contaminated with fragments from open chromatin regions. These stringent, high-salt conditions can potentially disrupt genuine biological interactions of interest, such as the association of factors that are more weakly bound to DNA. While the CUT&Tag approach is effective for profiling highly abundant and stably bound features of chromatin, such as histone modifications, it is limited in profiling less abundant and less stable complexes, including TFs and chromatin remodeling complexes. Importantly, no existing technology enables comprehensive genome-wide profiling of non-histone chromatin-associated factors in single cells, resulting in a critical gap in the single-cell toolkit in mapping TF and chromatin regulator binding across different cell types and physiological or disease contexts. To address these challenges, our Docking & Deamination followed by sequencing (D&D-seq) approach overcomes existing limitations by introducing a technology that records the presence of specific non-histone DNA binders directly in the DNA. This breakthrough allows us to profile chromatin-associated factors across a range of affinities, capturing both strong interactions as well as transient interactions that would otherwise be disrupted under high-salt conditions. Furthermore, D&D-seq integrates seamlessly into common single-cell workflows, supporting its broad adoption and enabling its integration with other molecular modalities for multi-omics profiling of gene regulatory networks at single-cell resolution. The D&D-seq approach provides significant technical and biological advances. Technically, it produces datasets that deliver accurate quantification of DNA:Protein interactions at single-cell resolution, contain sufficient information to identify cell-type-specific binding patterns, and match high-quality ChIP-seq data. For example, our cell line mixing experiment showed that D&D-seq successfully identified cell-specific TF binding patterns for CTCF, GATA1, recapitulating profiles generated using gold-standard protocols, such as ENCODE’s ChIP-seq. We further validated these results in primary human peripheral blood cells, demonstrating our protocol’s ability to profile the cell-type-specific binding pattern of non-histone DNA-binding proteins, such as CTCF, with high efficiency and accuracy. The richness of these datasets allowed us to apply predictive machine learning algorithms originally designed for bulk experiments to pseudo-bluked single-cell data, enabling the inference of chromatin 3D structure for each cell subpopulation. Crucially, D&D-seq opens new avenues for addressing fundamental biological questions that were previously inaccessible. For example, it can be applied to explore the dynamic regulation of chromatin remodeling complexes like SWI/SNF or Polycomb Repressive Complexes (PRC1 and PRC2) in heterogeneous cellular populations. Profiling these complexes at single-cell resolution is critical for understanding how they contribute to cell fate decisions during development or how they are dysregulated in diseases such as cancer. Similarly, D&D-seq can be used to study transient and context-dependent TF binding events, such as those of pioneer factors (e.g., FOXA1) that initiate chromatin opening or stress-responsive TFs (e.g., NRF2) that activate in response to environmental signals. Additionally, D&D-seq has the potential to uncover novel insights into epigenetic mechanisms in diseases, including neurodegenerative disorders and immune dysfunctions, where epigenetic rewiring is thought to play a role in pathology^48,49^. For example, profiling TFs and chromatin remodelers in single-cell populations of neurons or immune cells could reveal disease-specific regulatory circuits, enabling the identification of new therapeutic targets. Similarly, D&D-seq can be applied to study developmental disorders by mapping TF interactions during critical differentiation stages, where investigating interactions of factors such as SOX2 or NANOG in embryonic stem cells could shed light on the regulatory landscapes underlying pluripotency and differentiation. These applications highlight the transformative potential of single-cell profiling, as they enable researchers to untangle the intricate regulatory networks underpinning cell-type specificity, state transitions, and pathological alterations across different biological systems. Our work also demonstrates the power of integrating D&D-seq with single-cell genotyping. For example, we resolved the wild-type and mutant fractions of T cells in CHIP samples harboring an *IDH2* mutation. Using D&D-seq, we validated in vitro findings^40,41^ and directly observed aberrant CTCF binding profiles in mutant cells in vivo. Differential binding of CTCF at sites susceptible to *IDH2*-mediated changes in DNA methylation disrupted co-accessible regulatory regions and altered chromatin organization at critical loci, such as the *GIT1* (**Fig. 4k, 4l, 4m, 4n**) and *CNIH2* (**Supplementary Fig. 4h, 4i, 4j, 4k**) TADs. Interestingly, CD8 T cell-specific enhancers were found in both CTCF binding sites which were lost in the mutant T cells, suggesting dysregulation of enhancer interactions in this domain. In the mutant, enhancers at CTCF binding sites form a new co-accessibility with an upstream enhancer locus on NUFIP2, which is reported to regulate the expression of the inducible T-cell co-stimulatory receptor gene, ICOS^50^. ICOS plays an essential role in T cell proliferation and cytokine production in effector-memory T cells likely contributing to the elevated inflammatory state in CHIP^51^. Such differential enhancer regulation networks caused by methylation are well-studied in the imprinting of IGF2-H19, in which CTCF binding directs enhancers to H19 on the maternal allele while methylation at CTCF binding sites on the paternal allele redirects enhancers to IGF2, leading to distinct gene expressions^52^. This example underscores the potential of D&D-seq to uncover the molecular consequences of genetic mutations in human diseases and provide mechanistic insights into altered chromatin regulation. Notably, the flexibility of D&D-seq enables integration with other single-cell modalities, offering unparalleled opportunities for multi-omics analysis.

For instance, combining D&D-seq with CITE-seq or ASAP-seq allows simultaneous profiling of DNA-protein interactions and surface protein expression^53,54^, while integration with SHARE-seq, Paired-Tag, or CoTECH could link chromatin profiling with transcriptomic measurements^16,55,56^. This adaptability ensures compatibility with major single-cell technologies, such as the 10x Genomics Chromium system or split-and-pool approaches, expanding the range of biological questions that can be addressed relating to gene regulation across modalities, enabling analysis of gene regulation dynamics across the central dogma of DNA to RNA to protein^16,17^. Future integration of D&D-seq with tagmentation-based technologies, such as CUT&Tag^26^ or NTT-seq^24^, could enable profiling of DNA-protein interactions in regions with specific histone post-translational modifications. While our current work focused on integration with ATAC-seq due to the known enrichment of TF biding in open chromatin, integrating D&D-seq with single-cell whole-genome amplification methods (e.g., PTA, DLP+, DEFND-seq) holds promise for genome-wide mapping of DNA:Protein interactions^18,19,57^. These advancements will further expand the applicability of our tool, enabling researchers to tackle diverse questions in chromatin biology and gene regulation. A key limitation of D&D-seq is the relatively low number of editing events per cell. While our method allows for the first time to capture DNA:Protein interactions in single cells, the lower per-cell resolution makes it less suitable for studying individual cells with high granularity. The number of mapped binding events per cell may arise from antibody efficiency, circumvented in bulk methodology by using millions of cells as input. Despite this current limitation, D&D-seq excels in analyses that aggregate data across cells, such as pseudo-bulking or meta-cells^58^, which enable the discovery of complex biological patterns that were previously inaccessible. For example, the single-cell application allowed us to characterize a small subpopulation of CD8 T cells enriched in *IDH2* mutant cells in a primary CHIP sample. These cells exhibited a global reduction in CTCF binding, which perturbed chromatin architecture and reshaped the co-accessibility of specific topologically associated domains (TADs). However, we envision that future developments could significantly enhance the number of events per cell. For example, employing more active deamination enzymes with higher catalytic efficiency could amplify signal detection. Additionally, incorporating a secondary antibody staining step could boost the signal, allowing for a more comprehensive capture of DNA-protein interactions. These enhancements would further broaden the scope of D&D-seq, enabling its application to even more detailed and high-resolution studies at the single-cell level. In conclusion, D&D-seq represents a novel and versatile approach for multi-omics chromatin profiling, capable of measuring genome-wide distribution of non-histone chromatin-binding factors in bulk and single-cell samples. By tethering a DNA deaminase protein to specific sites in the genome, D&D-seq generates single-cell high-resolution DNA:Protein binding profiles with unprecedented accuracy. Its compatibility with diverse workflows and potential for integration with other modalities make it a transformative tool for addressing complex questions in chromatin biology. We anticipate that D&D-seq will empower discoveries across experimental systems and human tissues, providing new insights into DNA-protein interactions and their roles in health and disease.

## ACKNOWLEDGMENTS

We thank all members of the Landau lab for critical input. We thank Jane Park for outstanding technical support and Henry Rui for operational support. DAL is supported by the Burroughs Wellcome Fund Career Award for Medical Scientists, the Vallee Scholar Award, the Sontag Foundation (Distinguished Scientist Award, SFI 203261-01), the Leukemia Lymphoma Scholar Award and the Mark Foundation Emerging Leader Award. This work was supported by the National Heart Lung and Blood Institute (R01HL157387-01A1), National Cancer Institute (R33 CA267219), a Tri-Institutional Stem Cell Initiative award, the National Institutes of Health Common Fund Somatic Mosaicism Across Human Tissues (UG3NS132139-01) and the National Human Genome Research Institute, Center of Excellence in Genomic Science (RM1HG011014). This work was made possible by the MacMillan Family Foundation and the MacMillan Center for the Study of the Non-Coding Cancer Genome at the New York Genome Center.

## AUTHOR CONTRIBUTION

IR, W-YC, and DAL conceived the project and designed the study. W-YC, LM performed experiments. S-HY, W-YC, and IR performed bioinformatic analyses. W-YC, S-HY, SG, FI, DAL and IR analyzed and helped interpret the results. W-YC, S-HY, CP, DAL and IR wrote the manuscript. All authors reviewed and approved the manuscript.

## METHODS

### Cell culture

Human K562 (ATCC, CCL-243) and CA46 (ATCC, CRL-1648) cell lines were maintained according to standard procedures in RPMI-1640 (Thermo Fisher Scientific, 11-875-119) with 10% FBS (Thermo Fisher Scientific, 10-437-028) at 37 °C with 5% CO_2_. Cell lines in culture were screened biweekly for mycoplasma contamination using the MycoAlert PLUS Mycoplasma Detection Kit (Lonza, LT07-703).

### Primary cell acquisition and processing

Frozen PBMCs used for scD&D-seq were thawed into DMEM with 10% FBS, spun down at 4 °C for 5 minutes at 400*g* and washed twice with PBS with 2% BSA. Live cells from the PBMCs were enriched by the Dead Cell Removal Kit (Miltenyi, 130-090-101) per the manufacturer’s instructions.

### Cloning of nb-DddA constructs

Previously published sequences coding for the DddA11 enzyme were split at position 1397, and the C terminus of DddA11 (1290-1397) was synthesized as a gene fragment (Integrated DNA Technologies (IDT), **Supplementary Note**) flanked by restriction enzyme sites EcoRI and SpeI. The gene fragments were digested with EcoRI and SpeI for 1 hour at 37 °C, ligated for 1 hour at room temperature with pTXB1-alnbOc-Tn5 (Addgene, 184285), pTXB1-nbMmKappa-Tn5 (Addgene, 184286), or 3×Flag-pA-Tn5-Fl (Addgene, 124601) digested with the same enzymes. The ligated product was transformed into competent cells (New England Biolabs, C2987) per the manufacturer’s instruction. The final plasmid products (pTXB1-nb-DddA_NT) were confirmed by Sanger sequencing (Eton Bio) or Whole Plasmid Sequencing using Oxford Nanopore Technology with custom analysis and annotation (Plasmidsaurus).

### nb-DddA production

The pTXB1-nb-DddA_NT vectors were transformed into BL21(DE3)-competent *Escherichia coli* cells (NEB, C2527), and nb-DddA_NT was produced via intein purification with an affinity chitin-binding tag^24,59^. First, transformed bacterial colonies were grown overnight in 5 ml Luria broth (LB). Next day, 5 ml of LB culture was added to 400 ml and grown at 37 °C to optical density (OD600) = 0.6. nb-DddA expression was induced with 0.5 mM isopropyl-ß-D-thiogalactopyranoside (IPTG) and 50 µM ZnCl_2_ at 30 °C for 4 hours. After induction, cells were pelleted and then frozen at −80 °C overnight. Cells were then lysed by sonication in 30 ml HEGX (20 mM HEPES-KOH pH 7.5, 0.8 M NaCl, 10% glycerol, 0.2% Triton X-100) with a protease inhibitor cocktail (Roche, 04693132001). The lysate was pelleted at 10,000g for 20 minutes at 4 °C. The supernatant was transferred to a new tube, and 600 µl of neutralized 10% polyethyleneimine (Sigma-Aldrich, P3143) was added dropwise to the bacterial extract, gently mixed and centrifuged at 12,000*g* for 40 minutes at 4 °C to precipitate DNA. The supernatant was loaded on a 2-ml chitin column (NEB, S6651S), followed by washing with 12 ml of HEGX. Then, 3 ml of HEGX containing 100 mM DTT was added to the column with incubation for 48 hours at 4 °C to allow cleavage of nb-DddA from the intein tag. After incubation, 2 ml HEGX was added to elute nb-DddA directly into a 10-kDa molecular weight cutoff (MWCO) spin column (Thermo Scientific, 88527). Protein was dialyzed three times using 15 ml of 2× dialysis buffer (100 HEPES-KOH pH 7.2, 0.2 M NaCl, 0.2 mM EDTA, 2 mM DTT, 20% glycerol) and concentrated to 0.5-1 ml by centrifugation at 5,000 g. The protein concentrate was transferred to a new tube, mixed with an equal volume of 100% glycerol, and stored at −20 °C. After purification, the protein was denatured at 95 °C for 5 minutes, analyzed by SDS-PAGE gel (BIORAD, 4561085) and imaged by BIORAD ChemiDoc Touch Imaging System (**Supplementary Fig. 1b**).

### Deamination assay

DNA substrates, including lambda phage DNA (NEB, N3011) or 5’ 6-FAM-labeled 30-mer dsDNA oligonucleotides (IDT, **Supplementary Note**) were used for testing nb-DddA deamination activity. Deamination reactions of 250-1000 ng DNA were performed in 50 μl of deamination buffer (40 mM Tris-HCl pH 7.4, 50 mM KCl, 1 mM MgCl2, 1 mM dithiothreitol (DTT), 20 µM ZnCl2) with 50 µM nb-DddA_NT and 100 µM DddA_CT (Eton Bio, HPLC-purified, purity >95%, **Supplementary Note**). The reactions were incubated at 37 °C for 1 hour unless otherwise indicated. For lambda phage DNA, the products were then purified with 1.6X Ampure beads (Beckman Coulter, A63881), treated with 1 μl of T7 endonuclease (NEB, M0302) in 30 μl T7 buffer, further incubated at 37 °C for 15 minutes per the manufacturer’s instructions, and analyzed by an agarose gel (Invitrogen, A42135). For 6-FAM-labeled dsDNA oligonucleotides, 1 μl USER enzyme (NEB, M5508) was added to the deaminase-treated products, incubated at 37°C for 1 hour, denatured at 95°C for 5 minutes, and analyzed by 15% polyacrylamide TBE-Urea gel (BIORAD, 4566055). The cleaved DNA fragments were imaged by BIORAD ChemiDoc Touch Imaging System.

### Antibodies

Antibodies used were CTCF (1:100, Active Motif, 61932), GATA1 (1:100, Abcam, ab11852), and GATA2 (1:100, Invitrogen, PA1-100).

### D&D-seq

#### Cell fixation, Permeabilization, and Antibody binding

For all the following centrifugations, cells were centrifuged in a swing-bucket centrifuge at 300*g* for 5 minutes before fixation and 600*g* for 5 minutes after fixation. About 200K to 1 million cells were collected and resuspended in 400 µl PBS. Then, 16% methanol-free formaldehyde (Thermo Fisher Scientific, PI28906) was added for fixation (final concentration 0.1%) at room temperature for 5 minutes. Glycine (final concentration 125 mM) was added to stop the cross-linking, followed by a wash with 1 ml of PBS. Fixed cells were permeabilized in 200 µl permeabilization buffer (20 mM Tris-HCl pH 7.4, 150 mM NaCl, 3 mM MgCl_2_, 0.1% NP40, 0.1% Tween-20, 1% BSA, 1× protease inhibitors) on ice (7 minutes for cell lines, 5 minutes for primary cells), followed by a wash with 200 µl cold wash buffer (20 mM HEPES pH 7.6, 150 mM NaCl, 0.5 mM spermidine, 1% BSA, 1× protease inhibitors). After cell counting, 100K-400K cells were transferred to PCR tubes and resuspended in 100 µl antibody buffer (20 mM HEPES pH 7.6, 150 mM NaCl, 2 mM EDTA, 0.5 mM spermidine, 1% BSA, 1× protease inhibitors), with 1 µl (1:100 dilution) of the antibodies. The cells were incubated at 4 °C with slow rotation overnight.

### D&D binding and activation

The cells were washed once with 200 μl wash buffer, and another time with 200 μl D&D binding buffer (**Supplementary Note**). The cells were resuspended in 50 μl D&D activation buffer (**Supplementary Note**) with 100 uM DddA_NT, and incubated at room temperature on a rotator for 1 hour. The cells were then washed with 200 μl D&D binding buffer, resuspended in 50 μl DnD activation buffer (**Supplementary Note**), and incubated at 37 °C for 1 hour for deamination.

### Bulk ATAC

Tn5 adaptors were purchased from IDT. Adaptors (100 uM) (**Supplementary Note**) were annealed in TE buffer to form mosaic-end, double-stranded (MEDS) oligos by incubating at 95 °C for 5 minutes and then cooling at 0.2 °C per second to 12 °C. MEDS-A and MEDA-B were were mixed 1:1, and 2 µl was transferred to a new tube and mixed with 18 µl of TnY (in-house) enzyme after 1 hour at room temperature to allow for transposome assembly^60^.

After D&D activation, the cells were washed once with 200 μl 1x Tris-TD buffer (component) and resuspended in 50 μl 1x Tris-TD buffer with 9 μl loaded TnY (TnY volume depends on its concentration and needs to be titrated to get the optimal tagmentation). To initiate tagmentation, the reaction was incubated at 37 °C for 1 hour. To extract DNA, 1 μl 10% SDS, 2 μl proteinase K (NEB, P8107S), and 3 μl 0.5M EDTA were added to the reactions, which were then incubated at 55 °C for 1 hour, followed by column-based DNA purification per the manufacturer’s instruction (Zymo, D5205). For library preparation PCR, we used uracil-tolerant DNA polymerase (NEBNext Q5U Master Mix, NEB M0597S) and Nextera-compatible indexing primers (**Supplementary Note**) to amplify the purified DNA. To increase the efficiency of the initial gap fill-in step, we spiked in 1 μl of non-hot-start polymerase Bst 3.0 (NEB, M0374). The PCR steps are: **72°C for 5 min; 98°C for 30 s; 12 cycles of 98°C for 10 s, 55°C for 30 s and 72 °C for 30 s; followed by 72 °C for 5 min**.

### Single-cell ATAC

After D&D activation, the cells were resuspended in 1x nuclei buffer and counted using trypan blue and a Countess II FL Automated Cell Counter. For the remaining steps, we follow single-cell ATAC-seq per 10x protocol (version CG000209 Rev F, 10x Genomics) with the following modifications.

1. During the GEM generation and barcoding reaction (step 2.1), 2 μl of uracil-tolerant DNA polymerase (NEBNext Q5U Master Mix, NEB M0597S) was added to the barcoding reaction mixture to facilitate the first few rounds of PCR, where we expect to have some uridine in the reaction.

### Sequencing

The sequencing libraries were sequenced on a NovaSeq 6000, NovaSeq X, or NextSeq 2000 with dual indexed, paired-end 100 or 150-bp settings. Specifically, i5: 8 bp (16 bp for single-cell sequencing), i7: 8 bp, read1: 100 or 150 bp, read2: 100 or 150 bp.

### Genotyping of Targeted loci with Chromatin Accessibility (GoT–ChA)

We performed GoT–ChA according to the previously published paper with minor modifications^21^. Briefly, we performed whole cell fixation, permeabilization, antibody incubation, D&D binding and activation as aforementioned bulk D&D-seq. After D&D activation, the cells were resuspended in 1× diluted nucleus buffer (10x Genomics) and counted using trypan blue and a Countess II FL Automated Cell Counter. Afterwards, the cells were processed according to the Chromium Next GEM Single Cell ATAC Solution user guide (version CG000209 Rev F, 10x Genomics) with the following modifications:

1. During the GEM generation and barcoding reaction (step 2.1), 1 µl of 22.5 µM GoT–ChA primer mix was added to the barcoding reaction mixture. The primers used are IDH2 locus-specific primers **IDH2_R140_F1 (or IDH2_ R140_F2)** and **IDH2_R140_R1 (or IDH2_R140_R2)** (**Supplementary Note**). These primers allow for exponential amplification of the GoT–ChA fragments relative to the linear amplification of ATAC fragments. In addition, to facilitate the first few rounds of PCR, where we expect to have some uridine in the reaction, we spiked in 2 μl of uracil-tolerant DNA polymerase (NEBNext Q5U Master Mix, NEB M0597S) into the barcoding reaction mixture.
2. During the post-GEM incubation clean-up (step 3.2), 45.5 µl of elution solution I is used to elute material from SPRIselect beads. A total of 5 μl is used for GoT–ChA library construction, and the remaining 40 µl is used for ATAC library construction as indicated in the standard protocol.
3. To generate the GoT–ChA library, two additional PCRs were performed on the 5 µl set aside during step 3.2. The first PCR aims to amplify genotyping fragments before sample indexing and uses **P5** and **IDH2_R140_N1 (or IDH2_R140_N2)** primers (**Supplementary Note**) with the following thermocycler program: **95 °C for 3 min; 15 cycles of 95 °C for 20 s, 65 °C for 30 s and 72 °C for 20 s; followed by 72 °C for 5 min** and **ending with hold at 4 °C**. After a 1.2× SPRIselect clean-up, biotinylated PCR product is bound and isolated using Dynabeads M-280 Streptavidin magnetic beads (Thermo Fisher Scientific, 11206D). In brief, the beads are washed three times with 1× sodium chloride sodium phosphate-EDTA buffer (SSPE, VWR, VWRV0810-4L), added to the purified PCR product and incubated at room temperature for 15 minutes. The beads are then washed twice with 1× SSPE buffer and once with 10 mM Tris-HCl (pH 8.0) before resuspending in water. The bead-bound fragments are then amplified and sample indexed using **P5** and **RPI-X** primers (**Supplementary Note**) with the following thermocycler program: **95 °C for 3 min; 6–10 cycles of 95 °C for 20 s, 65 °C for 30 s and 72 °C for 20 s; followed by 72 °C for 5 min and ending with hold at 4 °C**.

Final libraries were quantified using the Qubit dsDNA HS Assay Kit (Thermo Fisher Scientific, Q32854) and the High Sensitivity DNA chip (Agilent Technologies, 5067-4626) run on a Bioanalyzer 2100 system (Agilent Technologies) and sequenced on a NovaSeq 6000 or NovaSeq X system at the Weill Cornell Medicine Genomics Resources Core Facility with the following parameters: paired-end 100 or 150 cycles; read 1N, 100 or 150 cycles; i7 index, 8 cycles; i5 index, 16 cycles; read 2N, 100 or 150 cycles. ATAC libraries were sequenced to a depth of 25,000-35,000 read pairs per cell and GoT–ChA libraries were sequenced to 5,000 read pairs per cell. A list of the primer sequences used in this study is provided in **Supplementary Note**.

### Cell sorting + Genotyping

Cryopreserved peripheral blood mononuclear cells from a CHIP donor with IDH2R140Q mutation were thawed and stained. Briefly, the cells were resuspended in staining buffer (BioLegend, 420201) and incubated with Human TruStain FcX (10 min at 4 °C; BioLegend, 422302) to block Fc receptor-mediated binding. Then, the cells were stained with CD8-BV650 (Biolegend, 344730), CD4-FITC (Biolegend, 344604), CD19-PE (BD biosciences, 561741), CD14-APC (Thermo Fisher, 47-0149-41), CD56-BV786 (BD biosciences, 564058) (1:100 for up to 106 cells in a final volume of 100 µl) for 20 minutes at 4 °C, and DAPI (Invitrogen, D1306). The samples were then sorted for DAPI− and single-marker positive cells (CD8 for CD8+ T cells, CD4 for CD4+ T cells, CD19 for B cells, CD14 for monocytes, CD56 for NK cells) using the BD FACSymphony™ S6 Cell Sorter **(Supplementary Note)**. The genomic DNA of the sorted cells was extracted using Puregene cell kit (QIAGEN, 158043) per the manufacturer’s instruction, and PCR-amplified using KAPA 2X mix (Roche, 07958927001) and IDH2_R140_F2 + IDH2_R140_R2 (**Supplementary Note**) primers. The linear PCR amplicon sequencing was performed by Plasmidsaurus using Oxford Nanopore Technology with custom analysis, and the genotype ratio was quantified by the number of reads mapped to IDH2 wild type and IDH2^R140Q^.

### Bulk Data analysis

Raw bulk D&D-seq FASTQ data were analyzed using FastQC for initial quality control^61^. Nextera transposase sequence and homopolymer G were removed using CutAdapt with “CTGTCTCTTATACACATCTCCGAGCCCACGAGAC” for R1 and “CTGTCTCTTATACACATCTGACGCTGCCGACGA” for R2 and “G{12}” parameters whenever identified in the FastQC report^62^. After trimming adapter and polyG sequences, processed FASTQ files were re-analyzed using FastQC and aligned to the Human reference genome (GRCh38) using BWA-MEM2^63^. Read alignments were subsequently sorted and indexed using samtools. To call D&D edits in bulk datasets, bam files were analyzed as described in the following *Extraction of D&D signal* section.

For TF motif discovery, genomic regions of 400 bp to 1 kb (indicated in the figures) centered at the peak summits were used to query the genome using MEME^64^. To visualize the genomic tracks on IGV, bigwig files were generated using the DeepTools bamCoverage function with the – normalizeUsing BPM option set^65^. ChIP-seq peak coordinates for CTCF, GATA1 and GATA2 for K562 cells were downloaded from ENCODE^20^.

### Single-cell data analysis

#### Preprocessing and annotation of 10x scATAC-seq data

Raw scATAC-seq datasets were aligned to the Human reference genome (GRCh38) and quantified using CellRanger-ATAC (version 2.1.0). The resulting peak/cell matrix was filtered for low quality cells and normalized using Signac (version 1.13.0)^66^. In detail, cells having 1) reads aligned to the blacklist region more than 0.05%, 2) less than 20% of reads aligned to peaks, 3) transcription start site enrichment score less than 3, 4) nucleosome signal higher than 4, 5) outliers having too many (average of peak counts + 2*standard deviation of peak counts) or fewer peaks (average of peak counts - 1*standard deviation of peak counts) were filtered out (**Supplementary Fig. 2b, 3a, 4b**). Multiplets were additionally annotated using AMULET and removed^67^. The preprocessed data were normalized by latent semantic indexing analysis, projected to low dimensions using uniform manifold approximation (UMAP), and clustered using the smart local moving (SLM) algorithm. Gene activity based on chromatin accessibility was inferred in order to annotate markers of each cluster. Finally, motif activity score was computed using ChromVAR with the JASPAR2020 core database for *Homo sapiens*^68,69^.

For the cell line data, eight clusters were defined with a clustering resolution of 0.6, and annotated as one of two cell lines based on the marker gene activity. Cluster 7, which was found in both K562 and CA46 clusters, was removed. The remaining number of cells was 5,840 cells (CA46, *n* = 4,108 and K562, *n* = 1,732). For PBMC data, 18 and 16 clusters were identified with a clustering resolution of 0.8 in replicate 1 and 2, respectively. Clusters were annotated based on marker gene activity and cell type prediction using Azimuth with a Human PBMC reference dataset^70,71^. For replicate 1, two clusters were additionally removed because two different cell types were mixed (cluster 7, CD8 T cell and monocyte) or it had overall low quality and lacked markers (cluster 13). The total of 5,413 cells in replicate 1 and 10,394 cells in replicate 2 remained.

### Genotyping of IDH2 mutation using GoT–ChA

Raw FASTQ files were first analyzed using FastQC to examine overall data quality and mutant allele at the targeted locus^61^. Based on a FastQC report, position of mutant allele, a read file with mutant allele, and primer sequence were identified. Input FASTQ files were then processed using the GoT–ChA analysis pipeline (https://github.com/landau-lab/Gotcha)^21^. In detail, raw data were first split into smaller files and quality filtering was performed with the “FastqFiltering” function. The R1 FASTQ file for replicate 1 and R3 FASTQ file for replicate 2 were used for genotyping with *c(37:39)* as a mutation site. After quality control of raw data, mutation state is annotated with the “BatchMutationCalling” function based on the provided wild-type/mutant and primer sequences. For both replicates, the primer sequence was set as “GATGGGCTCCCGGAAGACAGTCCCCCCCAGGATGTT”, and “CCG” and “CTG” were set as wild-type and mutant sequences, respectively. Results from each split data were merged into a single barcode and read count matrix using “MergeMutationCalling”. Genotype of each cell was assigned by manually setting thresholds based on distribution of log-transformed read counts (**Supplementary Fig. 4d**).

### Extraction of D&D signal

To summarize genome edits introduced by D&D in scATAC-seq data, a bam file produced by CellRanger was split into each cell type based on barcode sequence and cell annotation using sinto. For bulk ATAC-seq data, bam files aligned to the genome using BWA-MEM2 were analyzed^63^. After splitting bam files, the following five steps were performed to analyze D&D-mediated genomic variants (**Supplementary Fig. 2a**). All steps are integrated in the D&D analytic pipeline.

1. First, each bam file was preprocessed to remove uninformative and low quality read alignments. Duplicated reads were marked and simultaneously filtered using “picard MarkDuplicates” with a *REMOVE_ DUPLICATES=true* parameter. Read alignments with high mapping quality Phred score (>= 20), primary alignment, reads aligned to intact chromosomes, and those with properly aligned mates were retained using samtools (version 1.19).
2. Next, all single nucleotide variants (SNVs) found in each filtered bam file were collected using “bcftools mpileup” with following parameters, *-a FORMAT/AD, FORMAT/DP*,*INFO/AD --no-BAQ --min-MQ 1 --max-depth 8000*. The pileup result subsequently converted into the vcf format reporting SNVs supported by at least two reads supporting variants from minimum three aligned reads. These thresholds can be adjusted by users based on their data.
3. Germline mutations were then filtered based on loci and alleles from the gnomAD database and variant allele frequency higher than 10%. If available, custom databases in vcf format can be provided to additionally filter uninformative mutations.
4. Preprocessed bam files from step 1 were analyzed using MACS2 (version 2.2.9.1) to call peaks with *-f BAM -- nomodel* parameters. Peaks were then filtered with blacklist region annotation using bedtools (version 2.31.1)^72^. Motif analysis was performed using MEME Simple Enrichment Analysis (SEA)^73^ with HOmo sapiens COmprehensive MOdel COllection (HOCOMOCO)^31^ v11 core motif set to identify binding sites in peaks. Optionally, users can perform motif analysis using HOMER2 (https://doi.org/10.1038/s41586-024-07662-z) or a reference bed file generated by ChIP-seq.
5. Peaks harboring motifs of interest or overlapping with ChIP-seq reference tracks were classified as target peaks. Target peaks were resized to 200 bp (up/ downstream 100 bp from the motif center or peak summit with ChIP-seq) and overlaid with C-to-T and G-to-A variants identified in step 3. When multiple motif positions were found, a position with the highest motif score was chosen. Peaks without binding motifs of interest are classified as background peaks and were resized to 200 bp by taking +/− 100 bp from the peak summit.

### Evaluation of D&D signal

To compare D&D edit counts between target and background regions, edit counts per peak were summarized and signal-to-noise ratio (SNR) were calculated by dividing the number of C-to-T and G-to-A by the number of other variants. First, edits per peak were calculated by dividing SNV counts by the number of peaks in resized target and background peak regions. We assume that the frequencies of non-D&D edit (edits other than C-to-T) are consistent across the target and background regions, so this can be used to normalize the D&D edit counts. In this manner, SNR was then calculated by dividing the D&D edit counts by the mean of non-D&D edit counts. Footprint analysis for D&D edits was performed by counting the number of D&D edits in each base pair from randomly sampled target and background peaks, +-100 bp from the center of the motifs. The random sampling was repeated for 10 times and mean and standard deviation were used for visualization. For both the cell mixing experiment and primary blood cells, we used 200 randomly selected peaks. The number of subsampled peaks can be set by the user.

### Benchmark analysis with ultra-low-input cleavage under targets and release using nuclease (uliCUT&RUN)

Raw uliCUT&RUN data for CTCF and negative controls with no primary antibody were downloaded from GEO (GSE111121)^33^. The quality of raw FASTQ files was assessed using FastQC^61^ and the adapter sequence in both reads was trimmed using CutAdapt^62^ with *-a AGATCGGAAGAG -A AGATCGGAAGAG -m 21* parameters. Trimmed reads were aligned to the mouse reference genome (mm10) using BWA-MEM2 with 10 as the minimum seed length (*-k 10*)^63^. Read alignments with high mapping quality Phred score (>= 20), primary alignment, reads aligned to intact chromosomes, and those with properly aligned mates were retained from the resulting bam files using samtools (version 1.19). The processed bed file of CTCF ChIP-seq was downloaded from an independent study which used the same cell line (E14 mouse embryonic stem cells, GSE11431)^74^. Because the bed file was aligned to mm8, the genomic coordination was updated to mm10 using LiftOver^75^, sorted and merged for overlapping coordination using bedtools^72^, and modified to SAF format to use it as a custom reference for alignment. Processed bam files were aligned to the SAF using FeatureCounts (Subread package version 2.0.4) with *-p -- countReadPairs -F SAF* parameters^76^. Fraction of fragments in peaks was calculated by dividing the assigned fragment counts by the total fragment counts. The total of 4,105 CA46 cells with D&D edits from the cell mixing experiment were sampled to 10, 50, 500, and 5,000 cells with replacement and repeated for 100 times. K562 cells with D&D edits (*n* = 1,290) were downsampled to 10, 50, 500, and 1,000 cells in the same manner. To compare D&D to uliCUT&RUN, all reads harboring C-to-T or G-to-A tentative D&D edits with matched downsampled cell barcodes were collected. The fraction of fragments in peaks was calculated as described above using K562 CTCF ChIP-seq from ENCODE.

### Evaluation of CTCF binding in CD8 T cells

In our integrated PBMC dataset, CD8 and CD4 T cells formed a continuous cluster without a clear separation (**Fig. 4a**). Because the IDH2 mutation in these samples was specifically enriched in the CD8 T cell population (**Fig. 4b-d**), we re-clustered CD4 and CD8 T cell populations to extract high confident CD8 T cells and removed clusters expressing CD4 T cell or non-T cell markers and mixed CD4 and CD8 T cells. The remaining CD8 T cell cluster (*n* = 7,546) consisted of 2,132 (534 WT, 329 MUT, and 1,269 not genotyped) cells from replicate 1 and 5,414 (853 WT, 486 MUT, and 4,075 not genotyped) cells from replicate 2. These cells were further grouped into MUT-enriched (*n* = 6,437 cells) and WT-enriched (*n* = 1,109 cells) based on the clustering and genotype proportions (**Fig. 4h**). To compare differential CTCF binding between MUT and WT CD8 T cells, we leveraged the linear mixture model (LMM) from the GoT-ChA analytic pipeline which considers sample-specific batch effect and variability in cell numbers. LMM analysis was applied to 1,102 CTCF peaks harboring more than 5 D&D edits and found in more than 5 cells. We used a permissive false discovery rate (FDR) threshold of FDR ≤ 0.25, which identified 104 CTCF peaks with differential CTCF bindings between MUT and WT (**Fig. 4i**). Co-accessible peaks were identified using Cicero^46^. First, ATAC peaks in two replicates were merged using GenomicRanges::reduce and new count matrices were generated using Signac workflow for both replicates. Count data were loaded in R using the Monocle 3 workflow. Batch effect was normalized by aligning two replicates using *align_cds* with *preprocess_method = “LSI”, alignment_ group = “Dataset”* and *reduce_dimension* with *preprocess_ method = “Aligned”* parameters. Cicero object was generated with batch corrected CellDataSet and UMAP coordinates from Monocle 3 and co-accessible peaks were inferred using the *run_cicero* function as described in the Cicero workflow (https://cole-trapnell-lab.github.io/cicero-release/).

### Hi-C contact map predictions

To generate Hi-C contact map predictions, we used the pseudobulked ATAC reads and D&D reads for each cell type as the input for C.origami^35^. The bam files of ATAC-seq and D&D reads were first converted to bigwig files using DeepTools with the command bamCoverage -- normalizeUsing RPKM --binSize 1 --bam $bamfile -o $bigwig^65^. The bigwig file for D&D reads were then normalized against the bigwig of the ATAC and log2-transformed with a pseudocount of 1 using the command bigwigCompare --binSize 1 -b1 $DnD.bigwig -b2 $ATAC. bigwig -o $DnD.normalized.bigwig. The ATAC bigwig files and the normalized D&D bigwig files were used as the inputs for inference of HiC contact maps using C.origami. For predictions generated without incorporating CTCF binding information, the ATAC bigwig files, along with a bigwig file containing zeros for all regions, were used as input.

## Code Availability

Python and R scripts used in this study to analyze D&D-seq data are available on GitHub (https://github.com/sangho1130/DnD/). Detailed parameter settings and thresholds used in the analyses are described in the Methods. All analyses were performed using Python and R in a mamba virtual environment. Detailed software versions are also described in the Methods.

## Data Availability

Raw data and processed data files generated from cell lines will be made available at Gene Expression Omnibus (GEO). Processed data files generated from patient samples will be deposited at GEO. Patient raw sequencing data containing genomic sequences generated in this study will be deposited at the European Genome–Phenome Archive. The GRCh38 reference genome was used for alignment of single-cell ATAC–seq data (refdata-cellranger-atac-GRCh38-1.2.0), freely available from the 10x Genomics website (https://support.10xgenomics.com).

ChIP-seq datasets of CTCF (ENCFF111JKR), GATA1 (ENCFF844WTT), GATA2 (ENCFF997NUA) in K562 cell, human primary CD4 T cell (ENCSR470KCE), CD8 T cell (ENCSR116AKQ), B cell (ENCSR075NRV), monocyte (ENCSR162KZY), and NK cell (ENCSR856TKC); ATAC-seq dataset of K562 cell (ENCFF077FBI); Hi-C datasets of human primary CD4 T cell (ENCSR335JYP), CD8 T cell (ENCSR321BHC), B cell (ENCSR847RHU), monocyte (ENCSR236EYO), and NK cell (ENCSR971CJS) are from ENCODE.

---

**Supplementary Figure 1.**
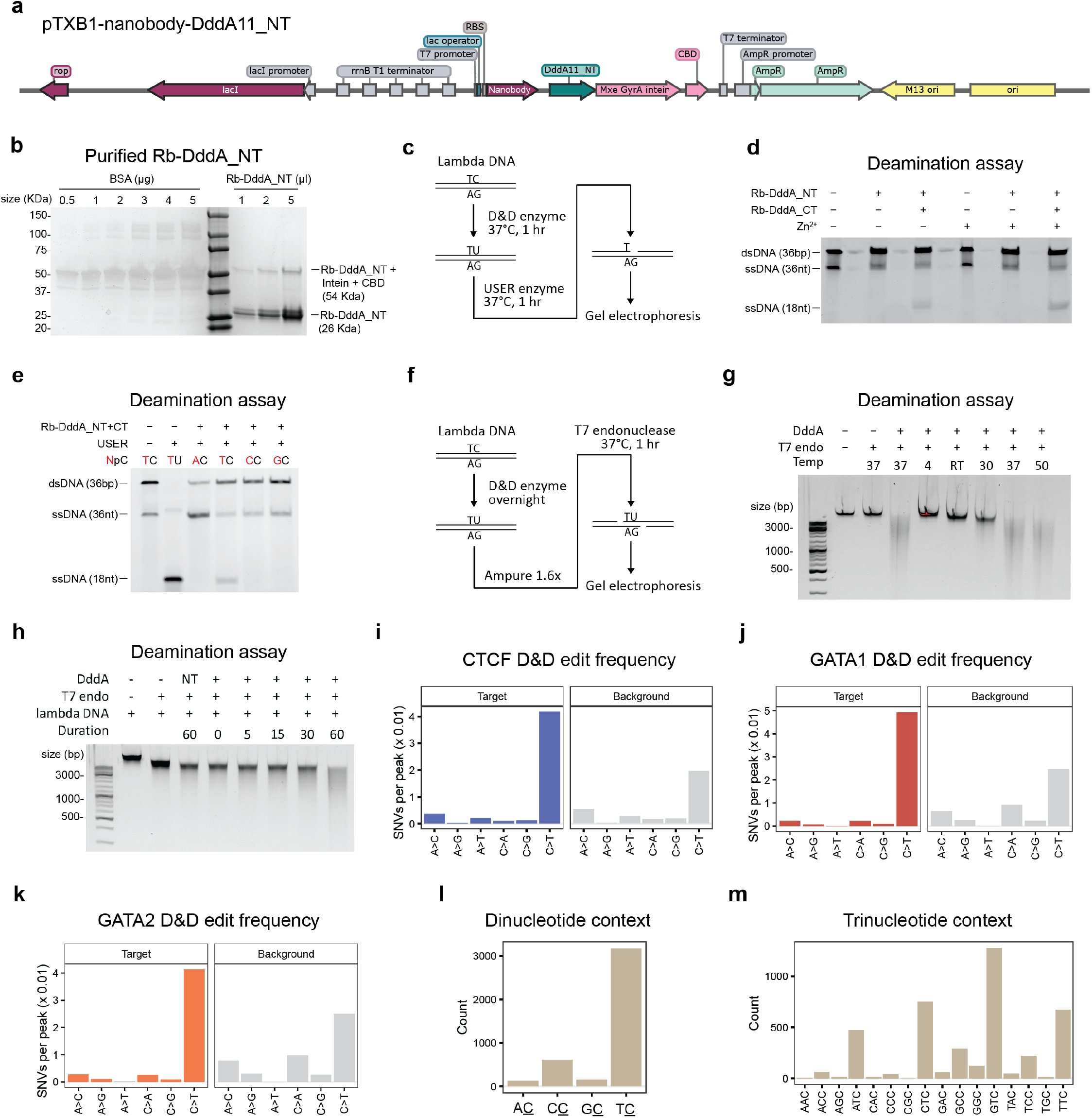
Docking & Deamination allow for mapping of transcription factor binding via molecular footprinting. **a)** Schematic representation of the pTXB1-nanobody-DddA11_NT plasmid. **b)** SDS-PAGE of purified Rb-DddA_NT protein. The upper band is uncleaved Rb-DddA_NT with intein and chitin-binding domain (CBD) (54 KDa). The lower band is cleaved Rb-DddA_NT (26 KDa). **c)** Scheme of in vitro deamination assay using 36-bp 6-carboxyfluorescein (FAM)-labeled DNA oligos treated with Rb-DddA_NT for conversion of C to U, followed by treatment with uracil-specific USER enzyme to create single-stranded nicks at U sites. Successful deamination can be detected by assessing single-stranded DNA, including generation of an 18 nt ssDNA fragment, on a denaturing gel. **d)** TBE-urea gel electrophoresis of DNA oligo treated with Rb-DddA_NT with or without Rb-DddA_CT and Zn2+. Rb-DddA_NT was activated upon addition of Rb-DddA_CT and Zn2+ further increased the activity. dsDNA, double-stranded DNA; ssDNA, single-stranded DNA. **e)** TBE-urea gel electrophoresis of DNA oligos containing different 5’ DNA contexts for the target cytosine treated with Rb-DddA_NT and Rb-DddA_CT. Rb-DddA preferentially deaminates cytosines that are preceded by thymines. **f)** Scheme of in vitro deamination assay using lamba phage DNA. **g)** E-gel of lamba phage DNA treated with Rb-DddA_NT+CT (DddA) overnight at different temperatures (°C). RT, room temperature. **h)** E-gel of lamba phage DNA treated with Rb-DddA_NT+CT (DddA) at 37°C for different duration (min). In lane 4, only Rb-DddA_NT is added as a negative control. **i)** DNA mutation signature of bulk K562 CTCF DnD-seq. Target regions are defined by ATAC peaks with CTCF motif (**Methods**). **j)** Same as (i) for GATA1. **k)** Same as (i) for GATA2. **l)** Dinucleotide context frequency of deaminated cytosine in K562 bulk CTCF DnD-seq data. **m)** Trinucleotide context frequency of deaminated cytosine in K562 bulk CTCF DnD-seq data.

**Supplementary Figure 2.**
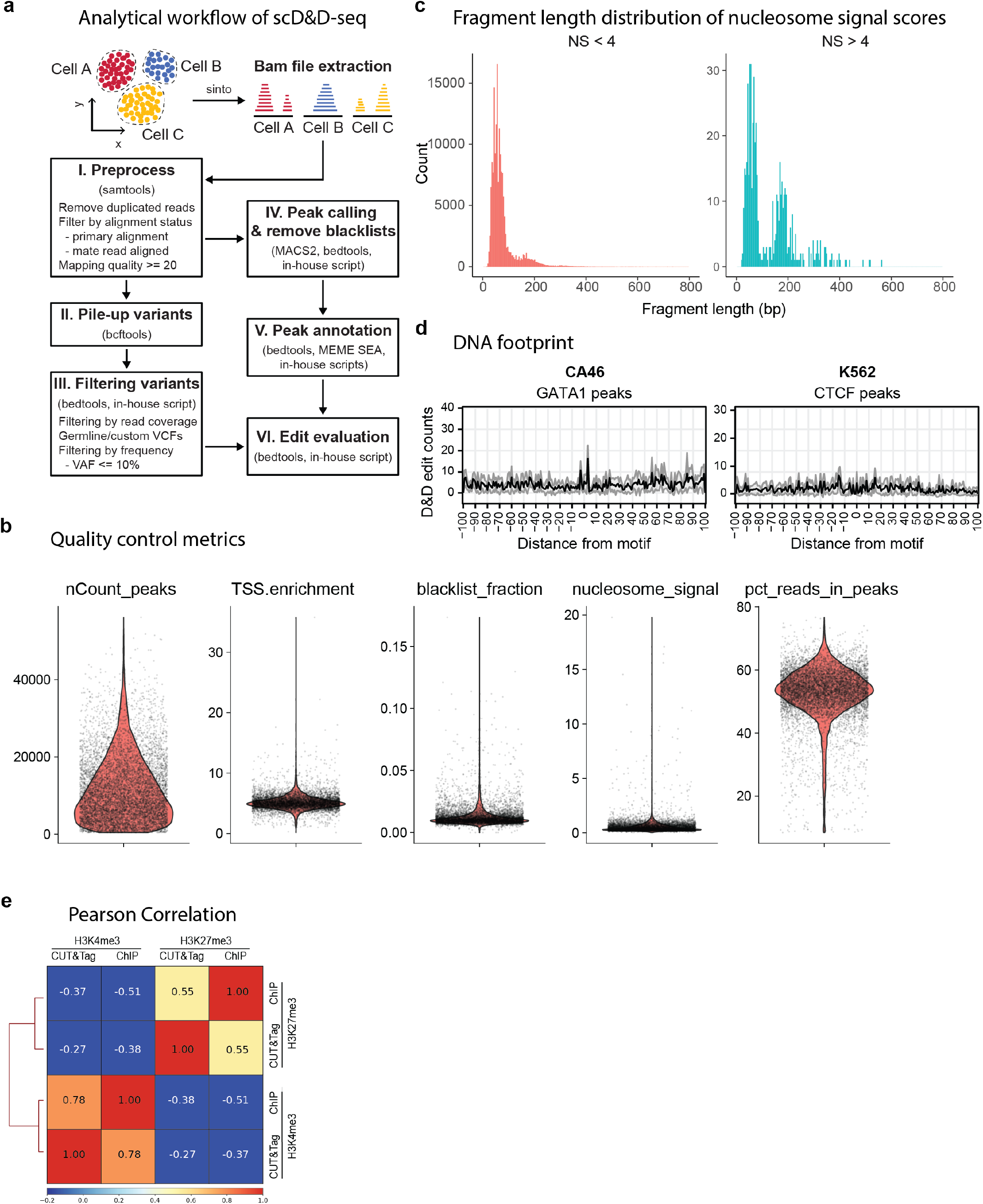
D&D-seq is compatible with existing droplet-based single-cell chromatin profiling technologies. **a)** Analytical workflow of scD&D-seq data. Reads from the same cell types were extracted from the bam file, followed by preprocessing and quality controls, variant pile-ups and filtering, peak calling, annotation, and finally evaluation of DnD reads. **b)** Quality control metrics of cell mixing scATAC-seq study, including number of counts in peaks (nCount_peaks), TSS enrichment scores (TSS.enrichment), fraction of reads in blacklist (blacklist_fraction), nucleosome signal scores (nucleosome_signal), and percentage of reads in peaks (pct_reads_in_peaks). **c)** Fragment length distribution of nucleosome signal scores (NS) <4 and > 4. **d)** Molecular footprint of pseudo-bulk D&D-seq for GATA1 in CA46 cells and CTCF in K562 cells showing count of C-to-T edits at aggregated sites showing no GATA1 binding signal in CA46 cells and no CTCF binding signal in K562 cells. **e)** Genome-wide Pearson correlation between CUT&Tag from ^26^ and ENCODE ChIP-seq for H3K27me3 and H3K4me3 in K562 cells. Peaks are defined by ENCODE ChIP-seq for H3K27me3 and H3K4me3.

**Supplementary Figure 3.**
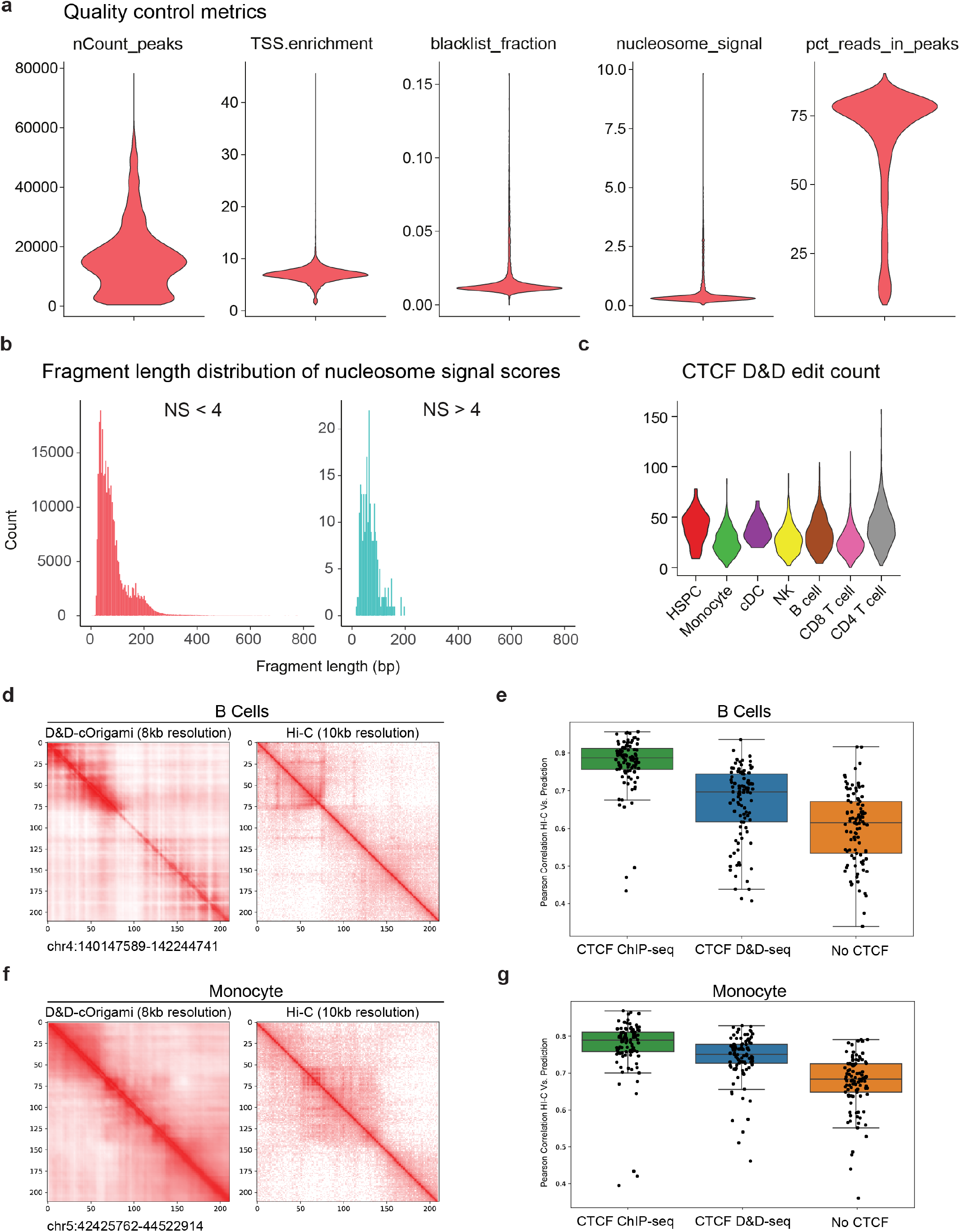
D&D-seq allows for DNA-protein interaction mapping in primary human cells. **a)** Quality control metrics of human primary PBMCs scATAC-seq, including number of counts in peaks (nCount_peaks), TSS enrichment scores (TSS.enrichment), fraction of reads in blacklist (blacklist_fraction), nucleosome signal scores (nucleosome_signal), and percentage of reads in peaks (pct_reads_in_peaks). **b)** Fragment length distribution of nucleosome signal scores (NS) <4 and > 4. **c)** Number of CTCF D&D edits per cell, separated by cell types. **d)** D&D-cOrigami-predicted (left) and experimental (right) Hi-C matrices of B cells. **e)** Pearson correlation of 100 randomly chosen B cells regions experimental Hi-C data (ENCODE) and: c.Origami 3D chromatin predictions generated with CTCF ChIP-seq data (ENCODE), CTCF D&D-seq or in the absence of CTCF information. Boxes represent the interquartile range, lines represent the mean, and error bars represent the 1.5 interquartile ranges of the lower and upper quartile. **f)** D&D-cOrigami-predicted (left) and experimental (right) Hi-C matrices of monocytes. **g)** Same as **e)** but for monocytes.

**Supplementary Figure 4.**
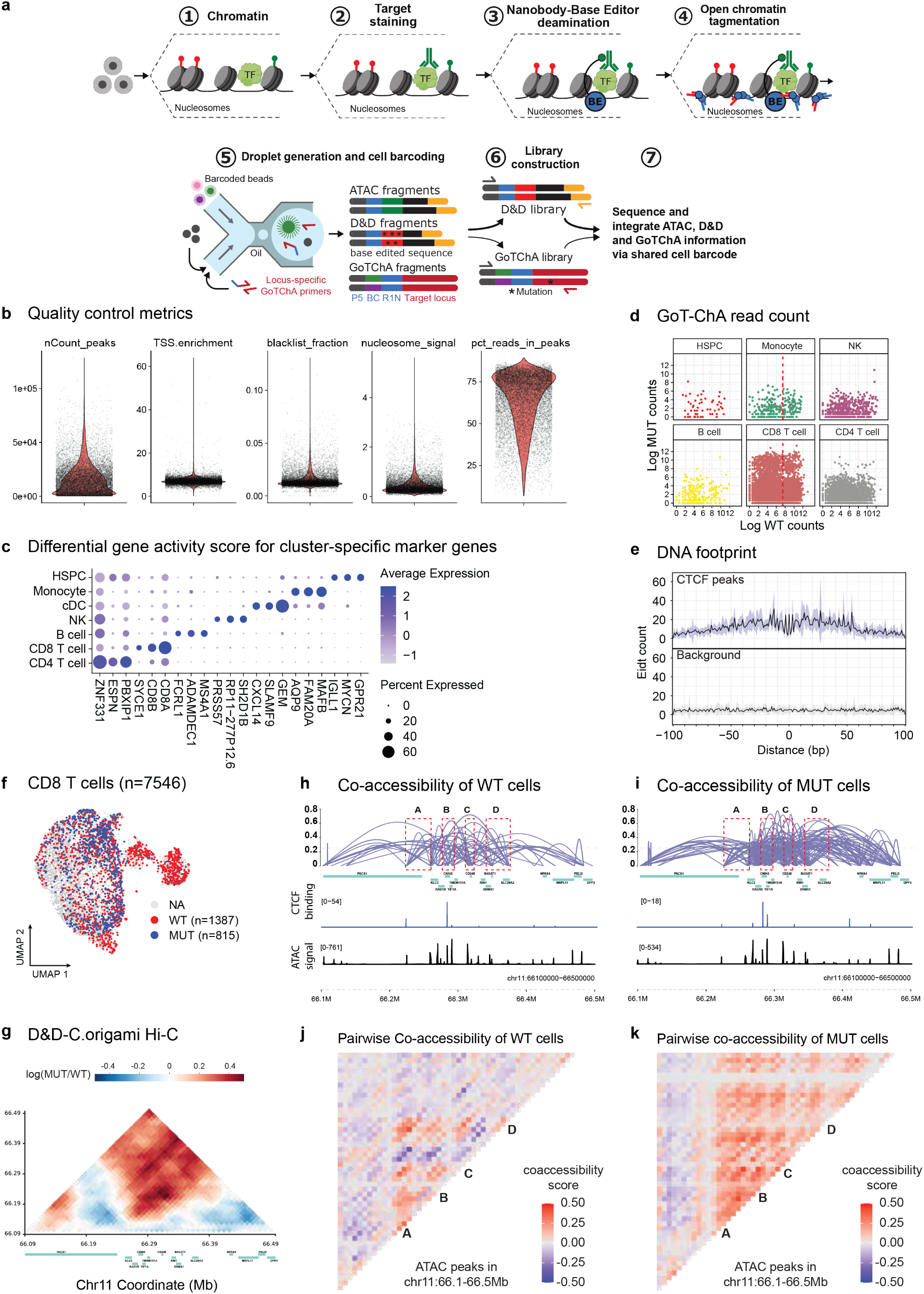
Multiomic application of D&D-seq to human primary clonal hematopoiesis samples. **a)** Schematic of D&D-GoT-ChA. **b)** Quality control metrics of scATAC-seq, including number of counts in peaks (nCount_peaks), TSS enrichment scores (TSS.enrichment), fraction of reads in blacklist (blacklist_fraction), nucleosome signal scores (nucleosome_signal), and percentage of reads in peaks (pct_reads_in_peaks). **c)** Differential gene activity score based on chromatin accessibility for cluster-specific marker genes. **d)** Distribution of WT and MUT read counts from GoT-ChA, separated by cell type. Thresholds are used to define MUT, WT and NA (Methods), with mutant cells in the top two quadrants. **e)** DNA footprint of on-target and background peaks. **f)** UMAP of the CD8 T cell fraction of the sample (n = 7,546 cells) with genotypes annotated. **g)** Heatmap displaying D&D C.Origami Hi-C predicted interactions enriched in mutant versus wild-type CD8 T cells at the genomic region on chr 11 encompassing the *CNIH2* locus. **h)** Co-accessible regions determined from ATAC data (purple lines) in wild-type CD8 T cells at the *CNIH2* locus. CTCF binding events and ATAC signal detected with D&D-seq are displayed in the bottom tracks. **i)** Co-accessible regions determined from ATAC data (purple lines) in mutant CD8 T cells at the *CNIH2* locus. CTCF binding events and ATAC signal detected with D&D-seq are displayed in the bottom tracks. **j)** Heatmap displaying pairwise ATAC-seq peak co-accessibility in wild-type CD8 T cells at the at the region corresponding to h and i co-accessibility plots. Co-accessibility score was smoothed by averaging three adjacent peaks. **k)** Heatmap displaying pairwise ATAC-seq peak co-accessibility in mutant CD8 T cells at at the at the region corresponding to h and i co-accessibility plots. Co-accessibility score was smoothed by averaging three adjacent peaks

